# Enhanced tissue penetration of antibodies through pressurized immunohistochemistry

**DOI:** 10.1101/2020.09.25.311936

**Authors:** Roberto Fiorelli, Gurpaul S. Sidhu, Arantxa Cebrián-Silla, Ernesto Luna Melendez, Shwetal Mehta, Jose M. Garcia-Verdugo, Nader Sanai

## Abstract

To address the inefficiency of passive diffusion for antibody penetration in thick tissue samples, which limits clearing-technique applications, we developed a versatile and simple device to perform antibody incubation under increased barometric pressure. Pressurized immunohistochemistry greatly improves the uniformity, intensity, and depth of fluorescent immunostaining in thick human and mouse brain samples. Furthermore, pressurized immunohistochemistry substantially decreases the time required for classic staining of thin sections.

**SUBMISSION CATEGORY:** New Results

## INTRODUCTION

Three-dimensional (3-D) histology is essential for exploring the mesoscale cellular architecture in brain tissue. Recently, the field was considerably advanced by the introduction of tissue-clearing techniques^1,2^ which resolve sample opacity generated by lipid-driven light scattering, allowing for 3-D imaging of rodent^3,4^ and human^5-9^ thick tissue sections (i.e., 100– 2000+ µm). Currently, researchers can choose among several clearing protocols,^1,2,5,10,11^ which differ by the equipment needed, chemicals required, procedure time, cost, and applicability. However, immunohistochemistry (IHC) on cleared tissue is challenging. Passive diffusion of antibodies is inefficient and does not generate uniformly deep stainings,^12,13^ limiting the use of clearing techniques, especially in human tissue. Proposed methods for enhanced antibody penetration using electric fields,^13^ system-wide binding controlling agents.^14^ and centrifugation or a peristaltic pump^15^ are difficult to reproduce,^7^ impractical to set-up, or are limited in the number of samples processed simultaneously.

In this study, we present an original experimental approach to IHC through the introduction of increased barometric pressure during antibody incubation. This experimental set-up does not require special equipment and allows simultaneous processing of multiple samples. Our findings demonstrate that pressurized IHC (pIHC) is a reproducible technique that greatly facilitates the speed and the depth of antibody penetration in cleared thick samples and thin brain tissue sections.

## RESULTS

Based on our experience in the preservation and IHC of postmortem human brain tissue,^16^ we aimed to adopt a clearing method suitable for these samples. We compared the performances of CUBIC (clear, unobstructed brain-imaging cocktails and computational analysis)^12^ and PACT (passive clarity technique)^17,18^ with slight modifications.^19^ Both techniques allowed 1-to 5-mm–thick samples of human caudate nucleus, cortex, and cerebellum to reach transparency in 14 days (Figure 1A). We noted tissue swelling of roughly 20% of volume with both procedures, as previously reported.^3,17,20^ Samples from human caudate nucleus reached up to 70% absolute transparency at the end of the procedure (Figure 1B). Samples richer in white matter, such as the corpus callosum (data not shown), required a longer time to reach such a degree of transparency compared with gray-matter-prevalent tissue. Preliminary clearing-coupled IHC tests using an indirect protocol failed to produce acceptable fluorescent signal deeper than 70 μm from the tissue surface, despite 72-hour incubations with primary and secondary antibodies.^3^

**Figure 1.**
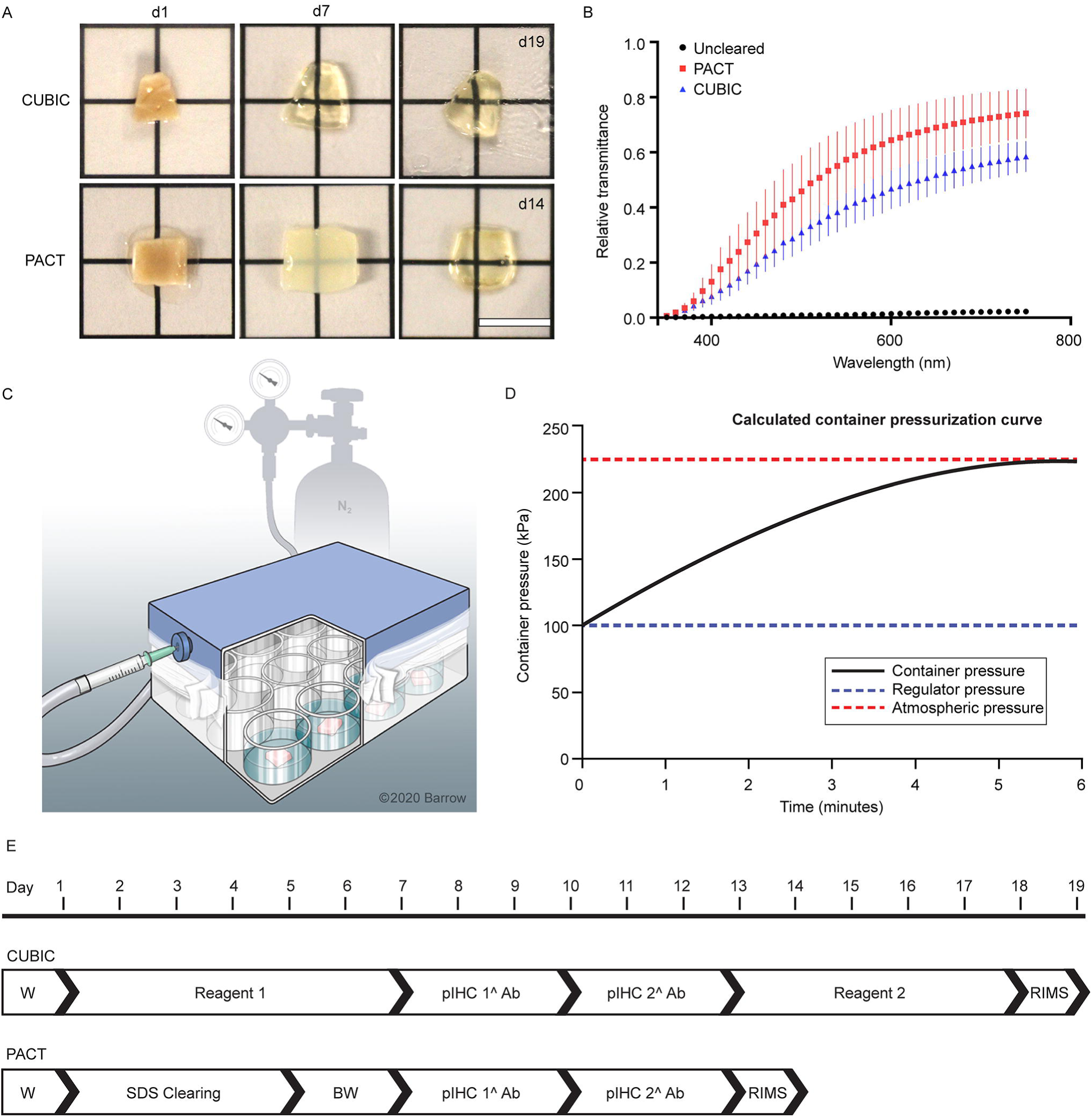
Pressurization apparatus and protocol. **(A)** Images of 1-mm–thick samples of human striatum during clearing procedures. Scale bar, 5 mm. **(B)** Graph showing the mean relative transmittance of the visible light spectrum measured in 10-nm intervals through striatal human samples. Black line: uncleared control sample. Red line: PACT-cleared sample. Blue line: CUBIC-cleared sample. Error bars show SE. N=3 independent samples per condition. **(C)** Schematic representation of the pressure device (see also Figure S1 for supplies). **(D)** Graph shows the calculation of changes over time in internal pressure in ideal conditions using a 225-kPa nitrogen gas inflow. Flow rate decreases exponentially because of the dynamic changes in the pressure differential between the enclosure and the gas source. **(E)** Comparative experimental timeline between CUBIC and PACT. Steps named Reagent 1, Reagent 2, and SDS Clearing summarize repeated incubations. Refer to Experimental Procedures for detailed protocol. 1^ Ab, primary antibody; 2^ Ab, secondary antibody; BW, boric acid wash; pIHC, pressurized immunohistochemistry; RIMS, refractive index matching solution; SDS, sodium dodecyl sulfate; W, washing step. *Used with permission from Barrow Neurological Institute, Phoenix, Arizona*.

### Increased Barometric Pressure Enhances Antibody Penetration

Thus, we hypothesized that antibody incubation in the presence of increased barometric pressure could improve the depth of the fluorescent signal. To test this, we built a simple airtight device (Figure 1C) using refurbished laboratory equipment (Figure S1A), wherein pressure was increased using N_2_ (Figure S1B, C and Video S1). With this set-up, 225 kPa was the safest maximum pressure allowed (Figure 1D). We then compared the intensity of clearing-coupled IHC when antibodies were incubated for 72 hours with pressurization versus free-diffusion conditions (Figure 1E). Technical details are available in the online Transparent Methods section of the Supplemental Information.

In a first test, antibodies for the microglia marker IBA1 (ionized calcium-binding adapter molecule 1), the marker for vascular smooth muscle cells, α-SMA (alpha–smooth muscle actin) and DAPI (4′,6-diamidino-2-phenylindole) were combined in a triple-channel IHC of human caudate nucleus samples. In free-diffusion conditions, CUBIC-treated samples demonstrated a tendency for higher permeability than PACT-treated samples (Figure 2A), evidenced by DAPI. Pressurized IHC significantly increased the intensity and depth of the staining for both clearing techniques compared to the controls (Figure 2B-E), with pIHC-CUBIC performing better than pIHC-PACT. IBA1+microglia were seen throughout the pIHC-CUBIC sample with no loss in morphological details (Figure 2E and Video S2). Our results exceeded depth reported by others for both IBA1^7,21^ and α-SMA.^9^

**Figure 2.**
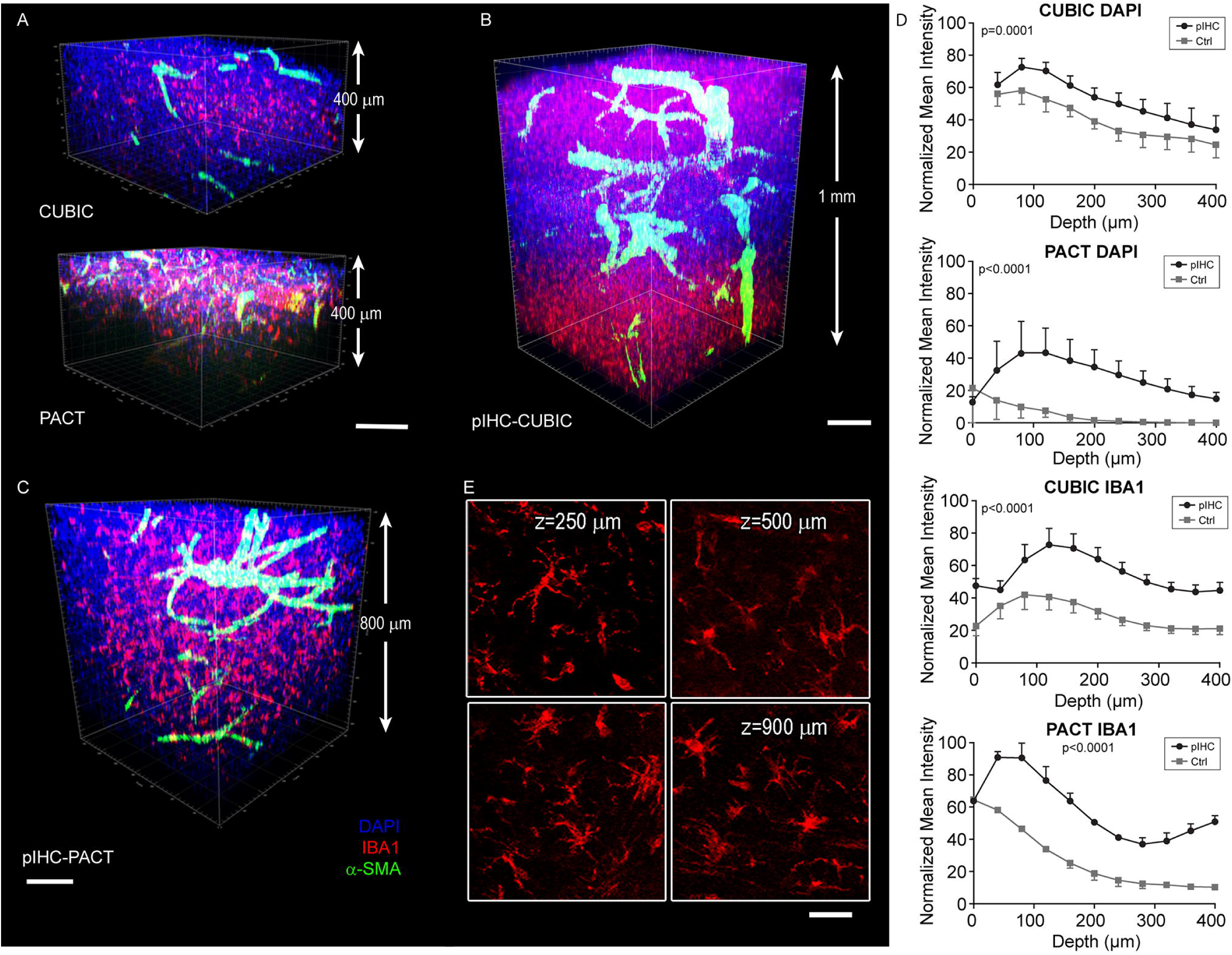
Pressurization increases penetration of microglia staining. **(A–C)** Cleared human striatal samples immunolabeled with DAPI (*blue*), IBA1 (*red*), and α-SMA (*green*). The 400-μm confocal acquisitions **(A)** show that free diffusion of antibodies generates uneven staining in both CUBIC and PACT-cleared samples. pIHC, independently of the clearing method (**B**, pIHC-CUBIC; **C**, pIHC-PACT), enhances the depth of antibody penetration. Images are z-stitched composites of 400-μm confocal acquisitions. Laser power was compensated for deeper acquisitions. **(D)** Graphs show quantification of normalized mean intensity measured along the z-depth at constant laser intensity and associated *P* values comparing pIHC with IHC. The x-axis shows depth from 0 μm to 400 μm. Pressurization (*black* line) significantly increases staining intensity compared to free diffusion (control, *gray* line), for both DAPI and IBA1 staining, independently of the clearing method. Error bars show SE. N=3 independent samples per condition. **(E)** Morphological details of IBA1+ microglia cells (*red*) are preserved across different z-depths in pIHC-CUBIC tissue. Scale bar in **A, B, C**, 200 μm; in **E**, 50 μm. All experiments were independently performed three or more times; representative images are shown. *Used with permission from Barrow Neurological Institute, Phoenix, Arizona*.

We tested pIHC with antibodies against GFAP (glial fibrillary associated protein), which are notoriously trapped by the astroglial network, impeding deep penetration.^22^ In cleared samples, pIHC led to more intense staining of astrocytes compared to controls, with pIHC-CUBIC again showing deeper staining than pIHC-PACT (Figure 3A-C and Video S3). However, pIHC-PACT revealed higher superficial intensity, lower background and superior details of GFAP+ processes (Figure 3D, E) compared with pIHC-CUBIC. In support of this observation, transmission electron microscopy (TEM) analysis revealed higher degradation of astrocytic intermediate filaments in CUBIC than PACT (Figure 3F).

**Figure 3.**
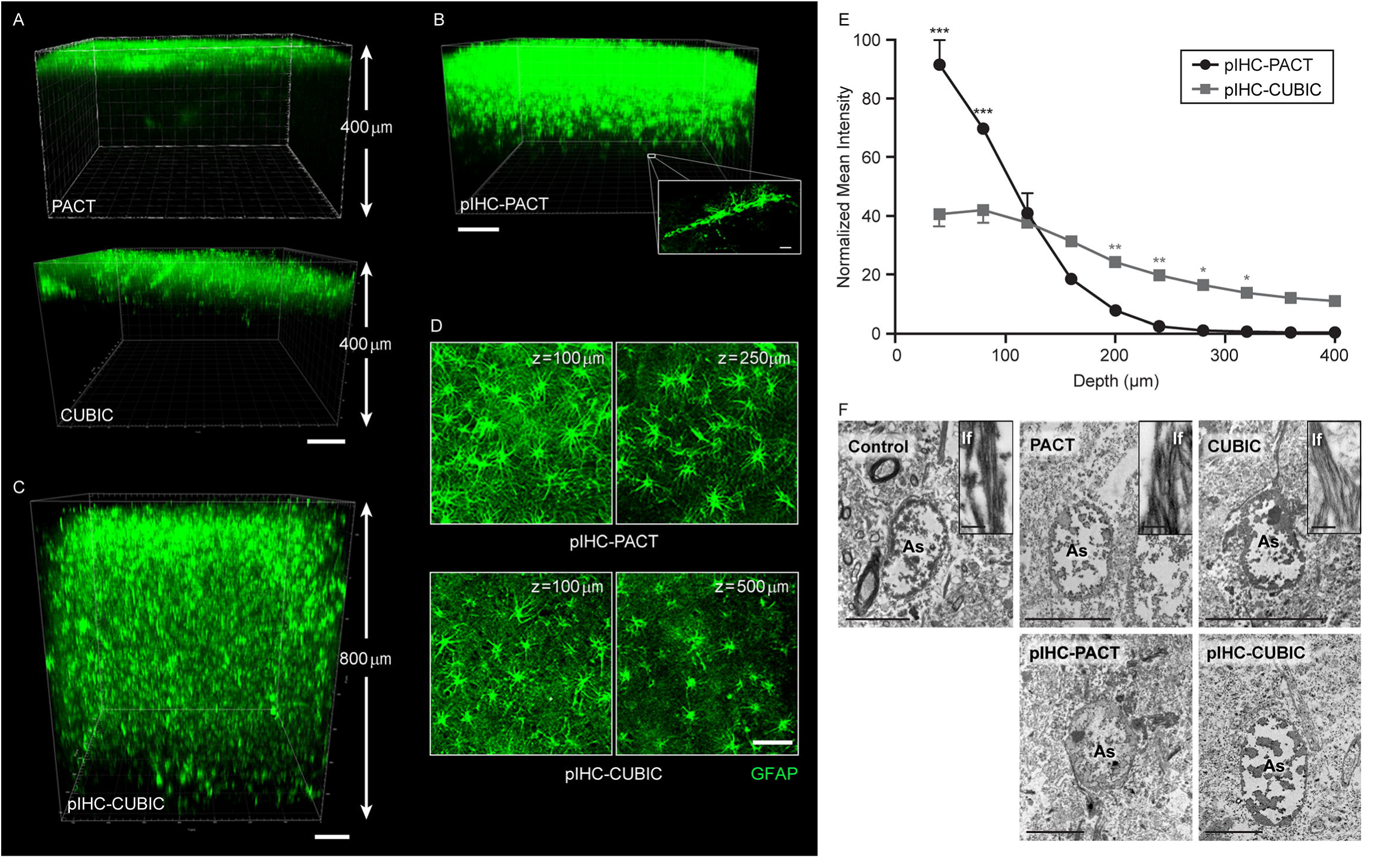
Pressurization increases GFAP penetration. **(A)** In PACT-cleared (top) and CUBIC-cleared (bottom) human striatal samples, GFAP (*green*) antibody is unable to effectively diffuse via traditional IHC. **(B)** pIHC enhances the depth of GFAP penetration in a PACT sample compared with free diffusion (compare with PACT in panel A, top image). Magnified box in B shows GFAP+ cells and astrocytic end-feet surrounding a blood vessel. These structures could be seen below maximum cellular staining depth. **(C)** pIHC for GFAP (*green*) in a CUBIC sample results in significantly deeper staining relative to free diffusion (compare with CUBIC in panel A, bottom image). Overall, GFAP staining in pIHC-CUBIC is deeper than pIHC-PACT (compare panel C and B). Image shows a z-stitched composite of 400-μm confocal acquisitions. **(D)** pIHC-PACT for GFAP (top) shows a superficial higher intensity and more detailed astrocytic processes than pIHC-CUBIC (bottom). Images show different z-depths. **(E)** Graph shows quantification of normalized mean intensity measured along the z-depth at constant laser intensity for pIHC-PACT and pIHC-CUBIC. pIHC-PACT (*black*) results in significantly more intense staining superficially (*black* asterisks) but quickly loses brightness with depth. Comparatively, pIHC-CUBIC (*gray*) demonstrates consistent intensity across depth, significantly reaching lower depths compared to pIHC-PACT (*grey* asterisks). Error bars show SE. N=3 independent samples per condition. **(F)** Transmission electron microscopy (TEM) micrographs of astrocytes (As) in different experimental conditions. Intermediate filaments (If) (in magnified boxes) are better conserved in PACT samples (middle column) than CUBIC samples (far right). Scale bar in **A, B**, and **C**, 100 μm; magnified box in **B**, 20 μm; in **D**, 50 μm; in **F**, 5 μm; insets in **F**, 250 nm. All experiments were independently performed three or more times; representative images are shown. *Figures 3A-E are used with permission from Barrow Neurological Institute, Phoenix, Arizona. Figure 3F is used with permission from University of València, València, Spain*.

We then tested pIHC on tissue cleared using the iDISCO technique. In line with the available protocol, samples were immunostained after the methanol-based pretreatment and permeabilization. The human striatal samples were labeled for IBA1, α-SMA, GFAP, NeuN (neuronal nuclei) and lectin (Figure S2). Imaging after the clearing procedure revealed that all stainings could be visualized in the superficial layers of the tissue, with depths between 50 and 100 µm. Despite the immunostainings lack of penetration, closer inspection of individual slices demonstrated their specificity and reliability (Figure S2A). In contrast to CUBIC and PACT, pressurization was not capable of increasing the immunostaining depth in iDISCO samples (Figure S2B, C). Therefore, we opted to focus on the characterization of the benefits of pIHC in CUBIC- and PACT-cleared tissue.

### Benefits of pIHC on Vascular and Neuronal Markers

Next, we stained CUBIC-cleared caudate samples for laminin, a ubiquitous marker of the vascular external lamina. pIHC, unlike free diffusion, led to intense and complete immunostaining of the human blood vessels throughout the tissue thickness (Figure 4A). Co-staining with α-SMA, which is expressed only in arterioles, showed that pIHC did not induce nonspecific staining and can be used for marker exclusion studies (Figure 4B). Furthermore, we performed pIHC on CUBIC-cleared human cortical samples with antibodies to the neuronal marker MAP2 (microtubule-associated protein 2) and the vascular-associated extracellular matrix marker fibronectin. Large samples (i.e., 5 mm × 5 mm × 5 mm) were used to preserve the intact pia membrane. Pressurization was again crucial to achieving deep staining because the pia membrane blocks antibodies (Figure 4C). All stainings could be imaged to a depth of 1500 µm (Figure 4D). Quantification of mean fluorescence intensity across the superficial 400 µm for laminin and fibronectin demonstrated a significant difference between control and pressurized conditions (Figure 4E). Similarly, pIHC for the vascular marker CD31 generated faithful immunolabeling throughout 1-mm striatal human samples (Figure S3A, B).

**Figure 4.**
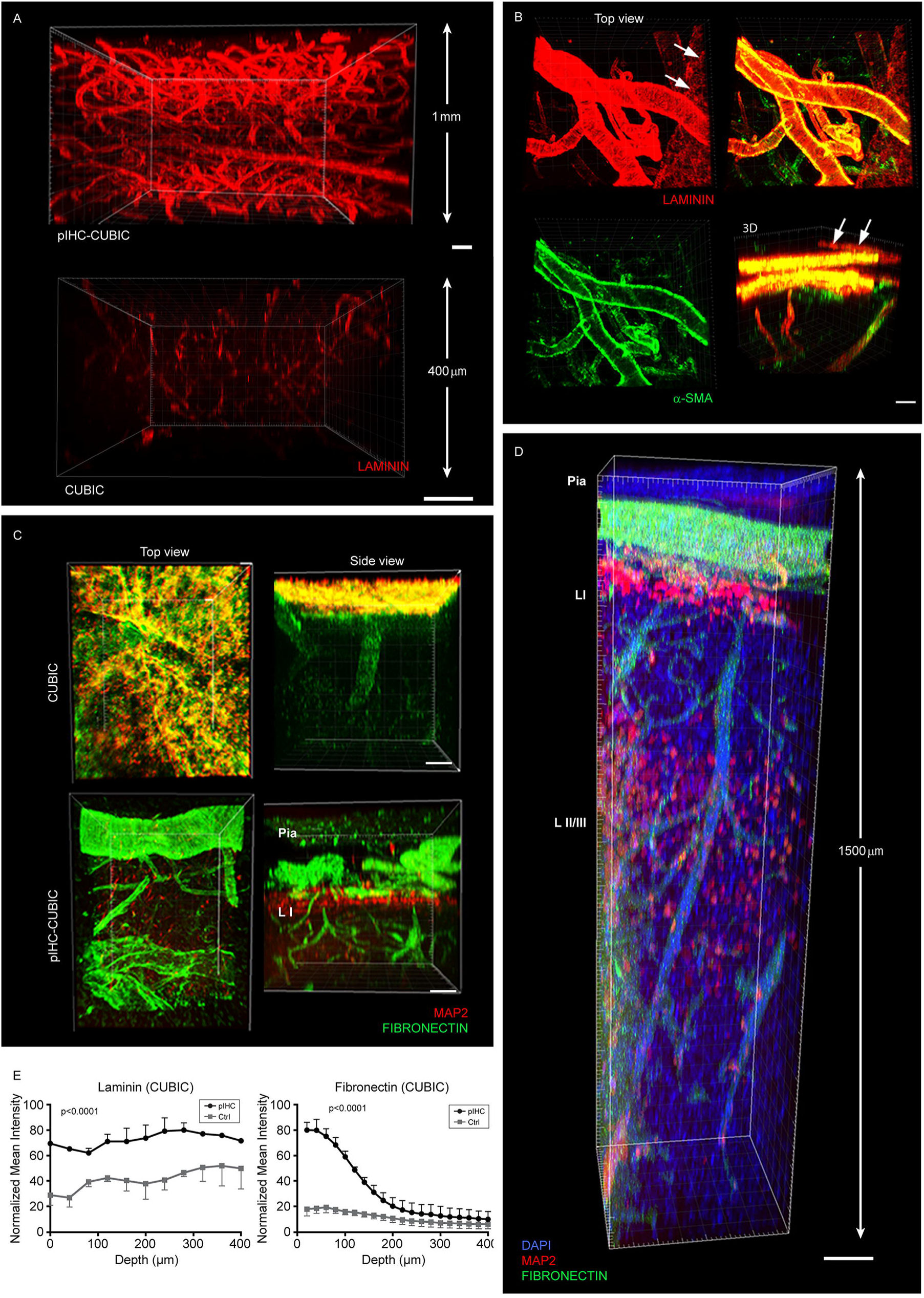
Benefits of pIHC in vascular and neuronal staining. **(A)** CUBIC-cleared human caudate nucleus tissue stained with pIHC (top image) for laminin (*red*) throughout the 1-mm z-plane. Free diffusion (bottom image) resulted in a dimmer and nonuniform staining pattern confined to the superficial 300 μm. **(B)** Two-dimensional (top and bottom left, top right) and 3-D (bottom right) images of pIHC-CUBIC–cleared cortex stained for laminin (*red*) and α-SMA (*green*) in pIHC-CUBIC–treated human cortical tissue. *Arrows* indicate a laminin+, α-SMA– negative blood vessel, while co-localization is demonstrated in remaining vasculature. **(C)** Top row: top (left) and side (right) views of CUBIC-cleared human cortex stained under free-diffusion conditions. The images demonstrate the pia membrane’s blockage of antibody penetration. Bottom row: top (left) and side (right) view of CUBIC-cleared human cortex immunostained using pIHC. Pressurization improves antibody penetration and reduces nonspecific fluorescence at the tissue surface. Side views show 400-μm z-acquisitions. **(D)** pIHC allowed for 1.5-mm–deep immunolabeling for fibronectin (*green*), MAP2 (*red*), and DAPI (*blue*) in CUBIC-treated human cortical sample. Fibronectin+ pial vessels perforate the cortex where cortical layer 1 (LI) and layer 2/3 (LII/LIII) are recognizable through the MAP2 staining. Laser power compensation was used to image the deeper tissue and reconstruct DAPI+/fibronectin+ blood vessels. **(E)** Graphs show quantification of normalized mean intensity measured along the z-axis at constant laser intensity and associated *P* values comparing pIHC with free diffusion (control, Ctrl) in CUBIC-cleared samples. Pressurization (*black* line) significantly increases staining intensity compared to free diffusion (Ctrl, *gray* line), for both laminin and fibronectin staining. In pIHC, the intensity of laminin remained consistent across z-depth while fibronectin intensity gradually decreased. Error bars show SE. N=4 independent samples per condition. pIHC-CUBIC, pressurized CUBIC; pIHC-PACT, pressurized PACT. Scale bars in **A, B**, and **D**, 100 μm; in **C**, 50 μm; all experiments were independently performed three or more times; representative images are shown. *Used with permission from Barrow Neurological Institute, Phoenix, Arizona*.

In contrast, pIHC of cerebellar neurons was challenging. Anti-neurofilament antibodies successfully stained processes of Purkinje cells throughout the molecular layer (Figure S3C, D), while the cellular dense granular layer blocked any staining below the superficial 120 µm. Two different anti-NeuN antibodies were tested with pIHC on CUBIC-cleared human hippocampus samples. Of these, only one (rabbit anti-NeuN from Cell Signaling) showed outstanding penetration that reached 400-µm depth even in free-diffusion conditions. Pressurization significantly increased the intensity of the signal in the mid-range z-depths, allowing reconstruction of the human dentate gyrus (Figure S3E-G). These observations add to the notion that the success of the staining is dependent on various factors, including the type of tissue and antibody used.

### Analysis of Effects of Clearing and Pressure on Tissue

We then conducted a whole-tissue deformation analysis to test for gross structural changes in the cleared tissue after pressurization (Figure S4). Interestingly, we found that CUBIC-cleared tissue dynamically changes size between different steps of the protocol (Figure S4A, B), settling for a final 25% increase compared to before treatment. Similarly, the size of PACT-cleared tissue oscillated during the protocol, finally showing a 50% increase when incubated in refractive index matching solution (RIMS) (Figure S4C, D). In both CUBIC and PACT testing, pressurization did not induce a significant variation in tissue size compared to control (Figure S4B, D).

Analysis of negative controls revealed that pressurization decreased tissue background independent of the clearing method (Figure S5A). Furthermore, significant autofluorescence of the entire vasculature was only observed in PACT (Figure S5A), while CUBIC erythrocytes lost their chromatic components, as shown by TEM and hematoxylin-eosin (H&E) analysis (Figure 5A and S5B). Additionally, PACT samples, unlike CUBIC, tended to accumulate non-specific granular deposits on the tissue surface (Figure S5C). TEM analysis revealed that pressurization caused an increase in protein aggregates in a network distribution (Figure 5B), which could relate to the decreased parenchymal autofluorescence observed. We observed no clearing-induced changes in the content of lipofuscin droplets (not shown), a typical source of autofluorescence in human postmortem tissue. H&E revealed a minor degree of vacuolization (Figure 5A) due to pressurization; however, cellular structures were not further disrupted, nor did pIHC enhance the artifacts caused by clearing-induced lipid degradation. Nuclear architecture in neurons could still be recognized, with partially intact chromatin (Figure 5C), while the rough endoplasmic reticulum was severely depleted in both treatments. Myelin components, neurofilaments, synaptic clefts, and plasma membrane were better conserved in CUBIC samples than in PACT samples (Figure 5D, E). Thus, with the exception of vacuolization, pressurization did not introduce additional artifacts beyond those already induced by each clearing protocol (compare CUBIC to pIHC-CUBIC, and PACT to pIHC-PACT, in Figure 5).

**Figure 5.**
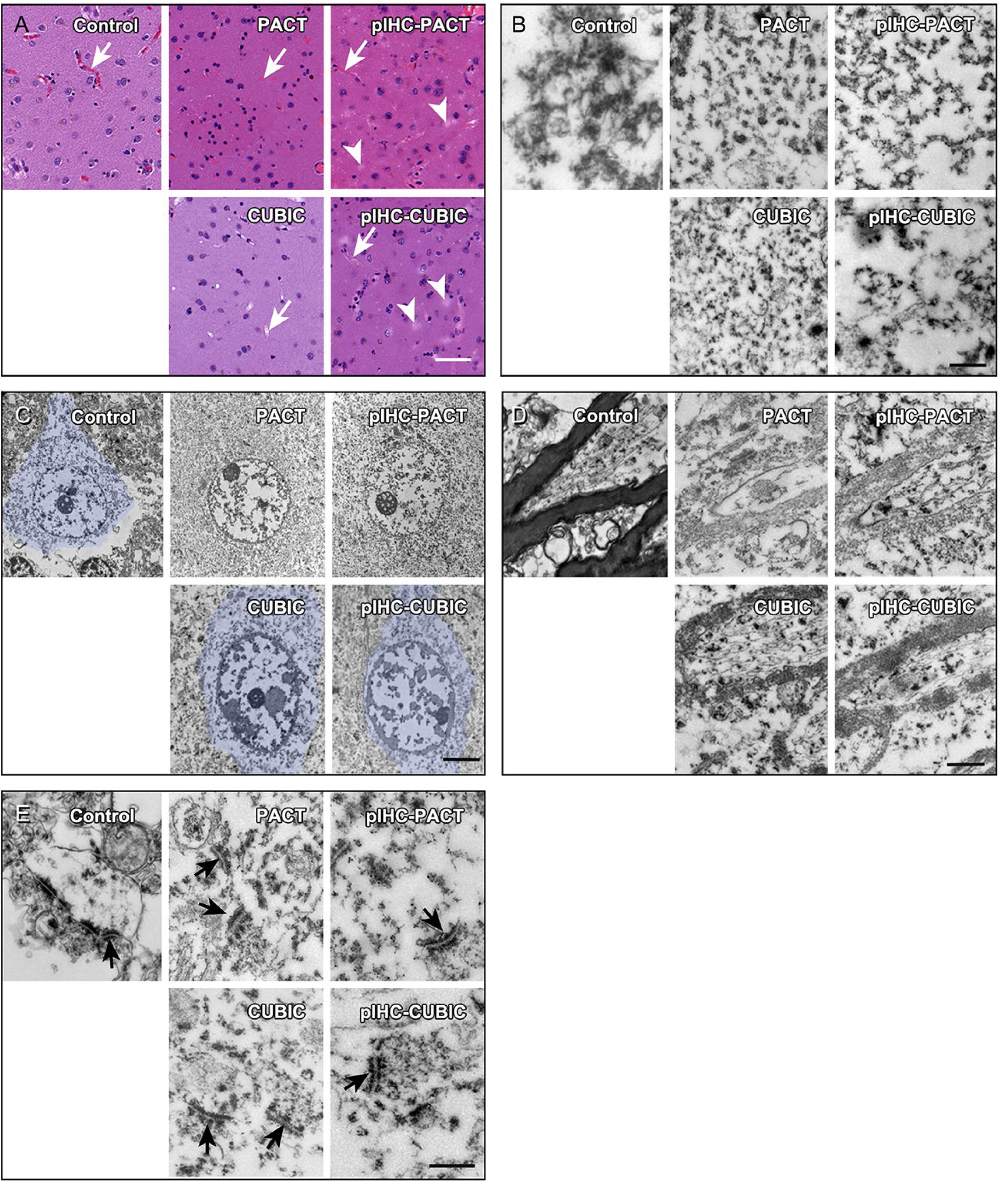
Analysis of treatment-induced artifacts. **(A)** Images show H&E staining of control, cleared (PACT, CUBIC), and cleared-pressurized (pIHC-PACT, pIHC-CUBIC) human caudate samples. Blinded examination by a pathologist revealed no discernable differences between the conditions. Some degree of tissue vacuolization (*arrowheads*) can be seen in the pressurized samples. Blood vessels (*arrows*) reveal erythrocytic discoloration in CUBIC but not in PACT samples and no differences in the respective pressurized samples. **(B)** High-magnification TEM micrographs of cellular cytoplasm show that pressurization (pIHC-CUBIC, pIHC-PACT) increases protein aggregates in a network distribution compared to unpressurized controls (CUBIC, PACT). **(C)** TEM micrographs showing the effect of treatments on cortical neuronal cells (pseudo-colored, *light blue*). The integrity of the cytoplasmic membrane was better conserved in CUBIC/pIHC-CUBIC than PACT/pIHC-PACT. **(D)** High-magnification TEM micrographs of myelin sheaths in different clarification methods of human cortical samples show better conservation in CUBIC/pIHC-CUBIC than PACT/pIHC-PACT. **(E)** Postsynaptic densities (*arrows*) are observed in all treatments with no difference after pressurization. The analysis was performed on 5 independent samples per each condition; representative images are shown. Scale bars in **A**, 50 μm; in **B**, 500 nm; in **C**, 5 μm; in **D** and **E**, 500 nm. *Figure 5A is used with permission from Barrow Neurological Institute, Phoenix, Arizona. Figures 5B-E are used with permission from University of València, València, Spain*.

### Validation of pIHC on Cleared Mouse Brain

Having validated pIHC-CUBIC in human tissue, we next tested it in a mouse model of human brain tumor, where a red fluorescent protein (RFP)-expressing primary human glioblastoma cell line (GB3-RFP) was injected unilaterally into the striatum (Figure 6A). Endogenous RFP fluorescence was preserved after clearing (Figure 6B). Pressurized incubation with tomato-lectin allowed staining of blood vessels across the entire tissue thickness (Figure 6C). Similarly, anti-vimentin antibodies revealed deep vessel-associated staining (Figure 6B) extending on the pial surface of the brain. Heat-induced antigen retrieval was adopted to immunostain CUBIC-cleared samples with the tumor stem cell marker SOX2 and the proliferation marker Ki-67 throughout tissue thickness. These markers were faithfully co-localizing with the RFP+ tumor environment and with migrating tumor cells along the corpus callosum (Figure 6D-E and Video S4). In tumor-free animals, we coupled pIHC with an anti-doublecortin antibody to detect neuroblasts along the neurogenic niche of the subventricular zone-rostral migratory stream (SVZ-RMS; Figure S6A-C).

**Figure 6.**
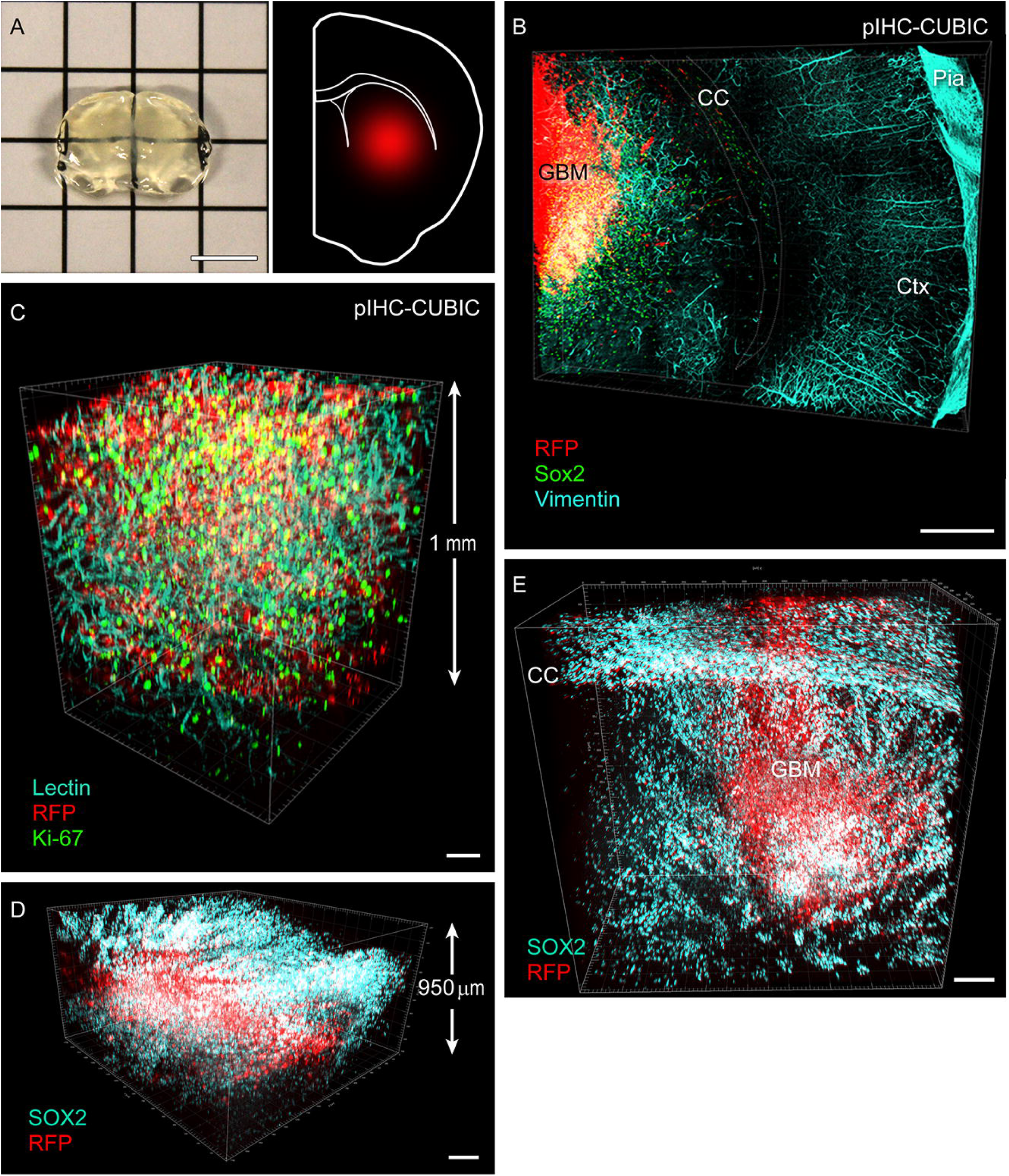
pIHC increases antibody penetration depth and speed in mouse tissue. (**A**) Left: photograph of 1-mm–thick mouse brain slice after CUBIC clearing. Right: schematic representation showing the location of the human glioblastoma xenograft (*red*) in a mouse brain hemisphere (*white*). **(B)** Mouse model of glioblastoma (GB3-RFP, red) stained with pIHC for SOX2 (*green*), and vimentin (*cyan*). Image shows a 400-μm–thick mosaic image of a 3 *×* 2.25-mm-wide mediolateral brain area. SOX2 co-localizes with the tumor core and migrating tumor cells. Vimentin shows a superficial pattern on the pial surface of the temporal cortex, together with a deep vascular pattern. **(C)** pIHC allows complete staining for Ki-67 (*green*), RFP (*red*), and tomato-lectin (*cyan*) through a 1-mm–thick tumor core area. **(D)** pIHC staining for the tumor marker SOX2 (*cyan*) across the entire section thickness. White cells show SOX2 co-localization with RFP+ tumor cells (*red*). **(E)** A 400-μm–thick mosaic confocal acquisition of a 1.8 *×* 1.7-mm-wide area shows striatal tumor core (GBM) with RFP+ (*red*) and SOX2+ (*cyan*) streams of migrating cells along striatal and callosal white matter traits. Scale bar in **A**, 5 mm; in **B**, 500 μm; in **C**, 100 μm; in **D**, 200 μm; in **E**, 200 μm. All experiments were independently performed three or more times; representative images are shown. CC, corpus callosum; Ctx, cortex; GBM, glioblastoma multiforme. *Used with permission from Barrow Neurological Institute, Phoenix, Arizona*.

### Pressurization Decreases the Time Necessary for Classic IHC Staining of Mouse Sections

Because pIHC improves antibody penetration, we also tested whether pIHC could achieve uniform staining in free-floating 40-µm–thick mouse brain sections faster than in free diffusion. The workload time was adjusted to perform the whole procedure in 8 hours with pIHC, compared to 19 hours for the standard protocol (Figure 7A). Tissue sections did not show damage due to pressurization. Tests with 8 different markers (i.e., GFAP, MAP2, IBA1, NF, OLIG2, Ki-67, NeuN, and tomato-lectin) showed no significant difference between pIHC and nonpressurized controls in the staining intensity across the z-plane, with the exception of IBA1, which showed a significant improvement under pressure (Figure 7B, C). As expected, the 8-hour procedure without pressurization yielded very weak signals for all 7 antibodies tested, and comparing it with pIHC further supports the role of pressure in facilitating antibody binding (Figure 7B). No differences were found in the lectin staining, for which 2 hours are typically sufficient to achieve complete staining in thin sections. This experiment shows that pIHC enhances the velocity of antibody penetration compared to free diffusion.

**Figure 7.**
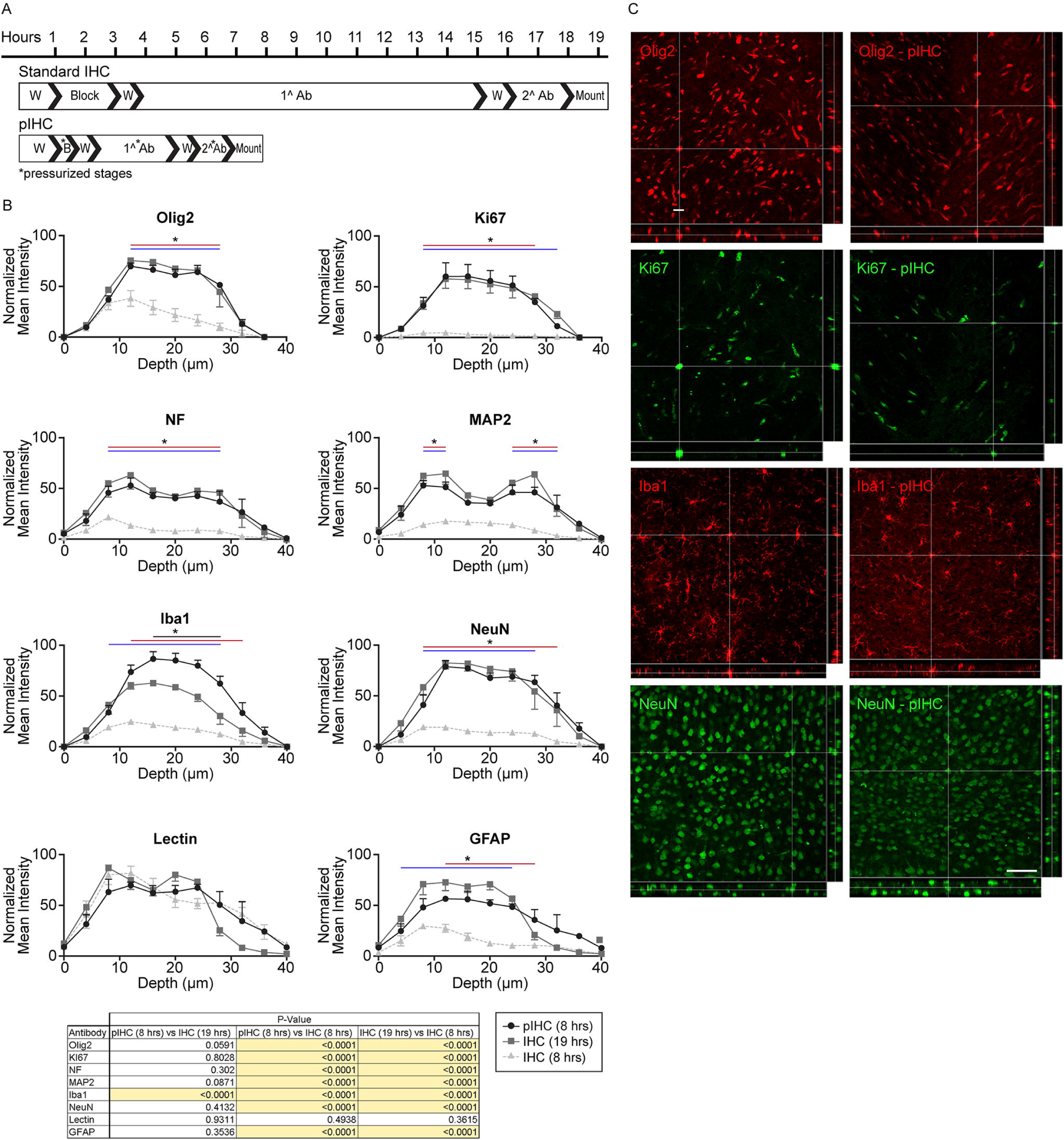
pIHC of 40-μm mouse brain sections. **(A)** Experimental timeline for classic (top) and pIHC (bottom) staining protocols on free-floating 40-μm–thick mouse brain sections. Pressurization (asterisk) was applied on blocking (B; standard: 2 hours; pIHC: 30 minutes), primary antibody (1^Ab; standard: 11 hours; pIHC: 3 hours) and secondary antibody (2^Ab; standard: 2 hours; pIHC: 1 hour) steps. Washing steps (W) were not subjected to pressurization **(B)**. Graphs show quantification of normalized mean fluorescence intensity across the z-depth and associated *P* values for a panel of 7 antibodies (OLIG2, Ki-67, neurofilament [NF], Map2, IBA1, GFAP) and one dye (tomato-lectin) tested on sections of mouse xenografts under three conditions: pIHC (8 hr), IHC (19 hr), and IHC (8 hr). pIHC (8 hr) (*black* line) shows no significant difference in staining intensity relative to traditional IHC (*gray solid* line), except for Iba1 that showed a stronger staining intensity with pIHC. Without pressurization, the 8-hour protocol (IHC 8 hr, *gray dotted* line) generated significantly weaker signal compared to pIHC (8 hr) and IHC (19 hr). The table shows the *P* values for all comparisons. N=4 independently repeated measurements. Error bars show SE. **(C)** Representative images of 4 immunostainings (OLIG2, Ki-67, IBA1, NeuN) performed with 19-hour standard protocol (left) and 8-hour pIHC (right) on thin mouse sections, revealed no differences in the staining intensity. Ki-67 and OLIG2 were imaged in tumor core areas while Iba-1 and NeuN were imaged in tumor-free cortical areas. A crosshair is used to indicate marker-positive cells located in the center of the section. Scale bar in **C**, 100 μm (all images). *Used with permission from Barrow Neurological Institute, Phoenix, Arizona*.

## DISCUSSION

This is the first time, to our knowledge, that an affordable and accessible method to improve antibody penetration in tissue via the use of a small increase in barometric pressure has been provided to the scientific community.

The technique of pIHC is based on a device of simple construction. All materials used are economical and available worldwide. Use of N_2_, an inert gas, does not introduce major artifacts in the tissue. For safety reasons, the pressure used did not exceed 225 KPa, which can be supported by a simple polyurethane box safely secured to a solid surface. Using pIHC, we achieved consistent and reproducible deep antibody penetration in thick tissue, and faster penetration in thin tissue compared to conventional free-diffusion incubation, while specificity was not altered. Importantly, pIHC was beneficial when coupled with aqueous-based de-lipidation clearing techniques, such as CUBIC and PACT, but not with the organic solvent-based iDISCO.

We hypothesize that the increased barometric pressure favors antibody diffusion through two concerted mechanisms. First, favorable diffusion conditions are formed immediately following initial pressurization. Because of tissue’s high hydrodynamic resistance, a temporary pressure gradient is created between the sample and its surroundings.^23^ This differential drives the antibodies into the tissue at an increased rate, relative to nonpressurized conditions, until pressure is equalized. Second, the increased atmospheric pressure generates an increase in the Brownian motion of the antibodies in solution, thus improving mixing. Because the box is a closed system, the added N_2_ mass during pressurization is trapped within a fixed volume. The overly abundant gas molecules more frequently enter the liquid surrounding the tissue, interacting with the antibodies within the solution. These collisions subsequently increase molecular motion and, as a result, improve the probability that the antibody will enter the tissue.

Future experiments are required to validate this mechanism empirically by comparing the sustained pressurization of pIHC to that of a pulsed protocol.^24^ By pulsing the pressure repeatedly throughout the incubation time, it would be possible to ascertain the extent to which the temporary pressure gradient affects antibody penetration. Further iterations of the pressurization device will allow live imaging of tissue during pIHC in order to determine the effect of increased pressure on mixing.

### Strategies for Enhanced Antibody Penetration

Despite the increasing number of approaches to tissue clearing being published every year, only a few methods have been proposed to improve antibody penetration, namely eTANGO.^13^ SWITCH (system-wide control of interaction time and kinetics of chemicals),^14^ and PRESTO (pressure-related efficient and stable transfer of macromolecules into organs.^15^ The eTANGO technique uses a rotating electrical field to force antibodies to penetrate the tissue and is compatible with both CUBIC and CLARITY but not iDISCO. Penetration in thick tissue of small labeling molecules such as SYTO-16 and lectin, as well as an anti-histone H3 antibody, is enhanced by electrotransport.^13^ Replicating this technique can be difficult as it requires advanced technology; however, devices capable of performing eTANGO have recently become commercially available.

The SWITCH approach solves the problem of antibodies being trapped on the most superficial layer of the tissue, by deactivating their binding sites using detergents. SWITCH is particularly interesting in histology because it allows immunostaining to be erased, permitting multiplexed imaging on a single sample. This useful application of SWITCH was validated with a large number of antibodies on medium-thin tissue sections (i.e., 100 µm thick). In thick tissue (i.e., 1 mm), the anti-histone H3 antibody was shown to effectively produce homogeneous, deep staining together with SYTO16 and lectin.^14^ Reproducibility of SWITCH is challenging as it requires specific clearing and immunostaining buffers, the composition of which is proprietary. Similar to eTANGO, SWITCH kits and devices are commercially available.

Two distinct approaches are possible with the PRESTO method, where antibody penetration is improved using centrifugal force (c-PRESTO) or by convection flow (s-PRESTO).^15^ Of the two PRESTO approaches, c-PRESTO is easy to reproduce because centrifuges are common in laboratories. Centrifugation induces a unidirectional force, generating a unilateral staining gradient, while unbound antibody is lost once it surpasses the tissue, therefore requiring repeated cycles of 12-hours of centrifugation with the addition of fresh antibody.^25^

In the second approach (i.e., s-PRESTO), the tissue is incubated with a large amount of antibody solution inside a syringe. The syringe plunger is pushed cyclically using a pump to create pressure, the extent of which is not explicitly stated. s-PRESTO allows for IHC to be performed on a limited number of samples in parallel. Importantly, the PRESTO techniques were shown to improve antibody penetration in extracellular matrix-rich peripheral tissues, but not in cleared rodent brain tissue. Using PRESTO, penetration of a NeuN antibody was improved in non-cleared rat brain tissue, but this improvement was limited to the superficial 100 µm.^15^

While pIHC shares the concept of applying pressure with s-PRESTO, there are fundamental differences in their application, the amount of antibody used, sample throughput, and accessibility. Overall, in the landscape of techniques for enhancing antibody penetration, pIHC features the innovative use of N_2_, a large breadth of tissue applications, ease of customization, and minimal costs for users.

Our structural analysis showed that pressurization does not affect the surface area of samples cleared with PACT, CUBIC, and iDISCO. Moreover, for PACT and CUBIC, the addition of pressure did not result in a statistically significant difference in surface area at every major step of the protocols. The consistency of these data strongly supports our assertion that pIHC does not significantly affect the macroscopic integrity of the tissue.

### Clearing of Human Brain Tissue

Investigation of the human brain tissue has shed new light on and identified phenomena specific to humans that cannot otherwise be observed in rodent models.^16,26^ Unlike the rodent brain, the human brain is cumbersome to study through fluorescent IHC because of the variability in procurement, fixation, preservation, and internal sources of autofluorescence. Postmortem human brain tissue can be difficult to obtain for ethical and practical reasons, especially in a short postmortem interval. Timely dissection and fixation are thus critical steps that can influence downstream applications such as IHC. Freshly extracted samples are most commonly fixed in a formalin or paraformaldehyde bath, while perfusion of human brain vasculature^15,27^ is exceptionally difficult to implement in a laboratory. Excessively long incubation in fixative is prone to over-fixing the tissue, masking epitopes from antibody recognition and thus making the tissue suboptimal for IHC. When treating tissue for confocal microscopy, 2-cm–thick fresh brain slabs reach optimal fixation after being submerged for 100 hours in 4% paraformaldehyde, as the speed of fixative penetration from both sides is 0.1 mm/h.^24^ Fixation can influence the time required for clearing, which for our samples was relatively short (i.e., 10 to 14 days). Notably, clearing time can increase up to several months in the case of long-term formalin-fixed^8,10,28^ or paraffin-embedded archival tissue.^7,9^

For the researcher who desires to adopt a clearing technique in the laboratory, the first step is to choose one of the many published methods. The type of tissue, desired clearing time, presence of endogenous fluorescent proteins, and compatibility with clearing-coupled IHC are parameters that must be taken into consideration (see comparative tables in the literature^5,10,11^). Particularly, when clearing valuable human specimens, it is imperative to consider the possibility of tissue damage. The use of strong solvents^29,30^ or electrophoresis^15,20,31^ can improve clearing speed at the risk of significantly deteriorating the tissue. Comparatively, CUBIC and PACT employ milder chemicals, which lower the risk of tissue damage^3,5,31,32^ but also increase the length of the procedure. The original CUBIC protocol was reported to have a lower efficiency in clearing human brain tissue than other organs.^9^ However, the application of a higher temperature^19^ (i.e., 40°C) during clearing allowed us to achieve acceptable transparency for CUBIC, as well as for PACT, with no significant differences.

### Factors Influencing Depth of Immunostaining

Examples in the literature show single immunostainings deeper than 500 µm in adult mouse or human brain samples, such as Arc,^33^ neurofilament,^10^ α-synuclein,^10^ tyrosine hydroxylase,^7,10,15^ IBA1,^34^ αβ-plaques,^11^ and parvalbumin.^35^ In most cases, images are limited to a single antibody while we demonstrate that pIHC allows for co-localization studies and quadruple stainings. In contrast, studies on embryonic tissue have reported successful deep imaging of large panels of IHC,^4,35,36^ due to the higher antibody permeability and inherent transparency of developing tissue compared to adult tissue.

Moreover, there is no consensus in the literature on the incubation time required to immunolabel cleared tissue, as it can range from 1 to 14 days.^7,15,21,30,34,36^ Therefore, when validating pIHC, we decided to utilize the incubation times suggested by the original CUBIC protocol, only modifying whether or not IHC was performed under pressure.

In our experiments, pressurization improved the depth of all markers used on human neural tissue, with a few exceptions. To increase the stringency of our protocol, we chose to perform pIHC incubations at 4°C, although temperatures in the range of 20° to 37°C can increase the depth of antibody binding^18^ and are the standard in cleared tissue staining.^4,7,9,10,21,30,34,35,37^ Notably, the CUBIC procedure involves subjecting immunostained tissue to a temperature of 40°C, which can cause minor fluorescence quenching.^19^ We also report that after long-term storage of immunostained samples in RIMS, the intensity was resistant to fading, contrary to other studies.^32^ When stored in darkness at 4°C, samples imaged 8 months after completion showed minor signal quenching.

As demonstrated by others,^10^ antibody diffusion was inversely proportional to the density of the target epitope, with GFAP being the most resilient to penetration while IBA1 antibody could stain throughout the tissue thickness. Although astrocytes and microglia are ubiquitous cell populations, there are obvious differences in the density of GFAP or IBA1 antigens. Accordingly, a neurofilament antibody could penetrate the cerebellar molecular layer, but not the granule cell layer (Figure S3C), the most cell-dense region of the brain.^38^ Conversely, the mouse SVZ-RMS region could successfully be stained for the neuroblast marker Doublecortin. The cell density in the RMS is similar to the cerebellar granule layer (respectively 2.7 million and 3.3 million cells/mm^3^)^38^; however, the success of this staining is due to the antibody bioavailability, which was significantly higher for the RMS given its small size. Other than areas of high cell density, the presence of extracellular-matrix-rich membrane can give rise to issues of surface adhesion, as in the example shown in Figure 4C. Here, pIHC was instrumental in allowing antibodies to overcome the pia membrane, and in producing an intense signal in the first layers of the cortex.

On the other hand, when employing vascular markers such as α-SMA, laminin, fibronectin, CD31, and lectin, pIHC consistently achieved complete staining through the tissue thickness. This result is due to two factors. First, vascular markers have a higher antibody bioavailability than cellular markers because the overall density of any vascular marker in each z-plane is generally lower than abundant cell populations. Second, pressurization likely facilitates a flow of the antibody solution through the vasculature. This phenomenon is suggested by the deep scattered GFAP-positive end-feet-enclosed structures shown in Figure 3A inset, which appeared in areas otherwise devoid of GFAP staining, suggesting that the antibody reached these areas through the vessels rather than the parenchyma. Similarly, vimentin antibodies revealed a vascular pattern in tumor xenograft samples (Figure 6B), while failing to penetrate beyond the surface in the tumor area because of antibody trapping by the vimentin network.

The success of particular immunostainings depends on the type of clearing because the integrity of antigens can be differently impacted by each clearing procedure, as suggested by the difference in staining quality of GFAP between pIHC-PACT and pIHC-CUBIC (Figure 3C). Because CUBIC degrades the intermediate filaments present in astrocytic networks, antibodies readily diffuse beyond the superficial layers of the tissue, resulting in deeper staining. Such degradation is possibly due to urea-dependent partial denaturation of proteins in CUBIC.^39^ On the other hand, recently developed delipidation-free and denaturant-free methods aimed to decrease the degree of tissue disruption,^7^ yet failed to produce IBA1 staining in contrast to the procedure chosen in our experiments.

In summary, differences in the depth and quality of clearing-coupled IHC can be attributed to tissue fixation, clearing method, tissue size, antigen density, and the antibody used.^10^ The empirical efficacy of all available antibody-diffusion enhancing techniques is dependent on the factors mentioned above. We have shown that pIHC greatly improves the depth and quality of the antibodies tested in the context of CUBIC and PACT. Future work will test the compatibility and efficacy of pIHC with alternative clearing methods as well as with tissues other than those in the brain.

## CONCLUSION

Overall, compared with conventional protocols, pIHC increases the antibody diffusion rate through tissue, allows for reproducible, deep immunostaining in thick samples, and significantly reduces protocol length in thin sections, without altering specificity. The device accompanying our approach can be constructed by any laboratory at a low cost compared to the considerably expensive devices, kits, or clearing-coupled IHC services offered commercially. pIHC could be of great interest for histology laboratories that are not yet using tissue clearing but are interested in speeding up the experimental turnaround of classic IHC. Further, the method offers great appeal for customization and application in molecular labeling techniques on any tissue thickness. Engineering of bench-top devices capable of reaching pressure levels higher than 225 kPa will provide the basis for further technique development. Implementation of pressurization can lead to a new generation of automated staining platforms, advancing clinical pathology with faster immuno-diagnostic protocols, which are highly desirable for real-time intraoperative diagnoses, particularly in surgical neuro-oncology.

## LIMITATIONS OF THE STUDY

There are several limitations to this study. First, empirical validation of the theoretical mechanisms requires considerable advancements in the apparatus design, to allow for real-time imaging of labeled particles and pressure pulsing. Second, in some cases, pIHC was not capable of overcoming the antibody network-trapping effect, e.g., vimentin staining in tumor tissue and neurofilament staining in the cerebellum. Since the identity of the antibody can directly affect its penetration capabilities, we advise users to test alternative antibodies against the same antigen or employ fragmented antibodies. Third, our experiments validate pIHC to achieve intense and homogeneous staining in 1-to 2-mm-thick tissue. Fully cleared brain IHC is the object of future validation of the technique. Additionally, pIHC is incompatible with the iDISCO technique, possibly due to tissue dehydration and lower delipidation compared to CUBIC and PACT. Future iterations of the device using higher pressures might solve this issue. Finally, vacuolization was observed in tissue samples subjected to pressurization. Although vacuolization did not affect the quality of immunostaining, we cannot exclude potential issues when attempting to visualize specific targets or structures. Therefore, when using pIHC, scientists should carefully consider the possible implications of vacuolization in their study.

## Supporting information

Permission letter - Fig 1, 2, 3A-E, 4, 5A, 6, 7, S1-S4, S5A, S5C, S6, and Vid S1-S4

Permission letter - Fig 3F, 5B-E, S5B

## DISCLOSURES

Figure 1, Figure S1, a, b, Figure 2, Video S2, Figure 3a, d, Video S3, Figure 5, c, Figure 4, a, b, c, d, Figure S5, a, b, Video S4, Figure 6, b, c, Figure 7, and Figure S3, were previously presented on July 17 2018, in a presentation entitled “Advancing 3-dimensional immunohistochemistry,” at the Barrow Neurological Institute Neuroscience Conference, Barrow Neurological Institute, Phoenix, AZ. Figure 1, Figure S1, a, b, Figure 2, Figure 3, Figure 5, Figure 4, Figure S5, a, b, Figure 6, and Figure 7 previously presented in a poster presentation titled “Enhanced tissue penetration of antibodies through pressurized immunohistochemistry,” at the Society for Neuroscience 2018 annual meeting, November 3-7, 2018, San Diego, CA.

## ACKNOWLEDGMENTS

The authors thank Dr. Tim Troxel, Dr. Jennifer Eschbacher, and Dr. Andrew Burnette for coordinating the human brain procurement and histopathological consulting, and Professor Mehdi Nikkhah and Professor Andrew Zydney for providing insights on the theoretical mechanisms. The authors thank the staff of Neuroscience Publications at Barrow Neurological Institute for assistance with the manuscript and video preparation.

## ABBREVIATIONS

α-SMA: alpha–smooth muscle actin;
c-PRESTO: centrifugal pressure-related efficient and stable transfer of macromolecules into organs;
CUBIC: clear, unobstructed brain-imaging cocktails and computational analysis;
DAPI: 4′,6-diamidino-2-phenylindole;
eTANGO: electrostochastic transport of activity modulated molecules in nanoporous gel organ hybrid;
GB3: glioblastoma 3 cell line;
GFAP: glial fibrillary associated protein;
H&E: hematoxylin and eosin;
IBA1: ionized calcium-binding adapter molecule 1;
IHC: immunohistochemistry;
MAP2: microtubule associated protein 2;
NeuN: neuronal nuclei; NF, neurofilament;
PACT: passive clarity technique;
PB: phosphate buffer;
PBS: phosphate buffered saline;
PBTX: PB/0.1% Triton-X;
pIHC: pressurized immunohistochemistry;
PRESTO: pressure-related efficient and stable transfer of macromolecules into organs;
R1/2: reagent 1/2;
RFP: red fluorescent protein;
RIMS: refractive index matching solution;
s-PRESTO: convection flow-related efficient and stable transfer of macromolecules into organs;
SVZ-RMS: subventricular zone-rostral migratory stream;
SWITCH: system-wide control of interaction time and kinetics of chemicals;
TEM: transmission electron microscopy;
2-D: two-dimensional;
3-D: three-dimensional

