## Supplemental Methods for "Enhanced tissue penetration of antibodies through pressurized immunohistochemistry"

### SUPPLEMENTAL METHODS: TRANSPARENT METHODS

#### Pressure Device

A polypropylene box (VWR International, Radnor, PA; dimensions of the internal chamber: 6 cm × 11.8 cm × 8 cm) was drilled, equipped with a 1.5-cm–diameter rubber stopper (BD Biosciences, Franklin Lakes, NJ), which was sealed with rubber cement. A 24-well plate (Corning, Corning, NY) was sawed to obtain a 12-well plate that could be secured in the pressure box (Figures 1C, S1A, and Video S1). As the tissue was submerged in staining incubation solution inside the multiwell (Figure 1C), the device was thoroughly wrapped with three layers of paraffin film (Parafilm, Bemis Company, Inc., Neenah, WI) and adhesive tape (3M, Maplewood, MN) for airtightness (Figure S1B) before proceeding to pressurization.

Tanked N_2_ equipped with a multistage gas regulator (VWR, cat. no. 55850-424) and two-stage manometers was employed as the gas source. The maximum pressure that could be passed through the needle was determined to be 225 kPa. Before pressurization, the box, the needle, and the inlet tube were securely immobilized for safety (Figure S1C). Using a 21-gauge needle, N_2_ was injected into the box for 10 minutes with an initial mass flow rate of 2.1 ± 0.1 mg/s (i.e., a gauge pressure of 125 kPa). Swelling of the paraffin film sealing (Figure S1B and Video S1) was considered as an indicator of pressure over time.

Because of equilibration between the chamber, hose, and the regulator pressure differential, pressure within the box could not exceed the inlet limit. To ensure that the inlet time was sufficient to theoretically reach that pressure, we utilized the orifice mass flow rate equation in combination with the ideal gas law to determine the time required for pressurization.

To calculate the flow rate of N_2_ into the container, we employed the following equation (1)^40^:

$$Q_{m}=\frac{C_{d}}{\sqrt{1-\beta^{4}}}*\epsilon*A_{o}\sqrt{2*\rho_{u}*\Delta P}$$

(1)

where *Q_m_* is the N_2_ mass flow rate through the needle, *C_d_* is the coefficient of discharge, *β* is the ratio between the diameter of the orifice and the diameter of the pipe, *A_o_* is the cross-sectional area of the orifice, *ρ_u_* is the upstream density of nitrogen, and *∆P* is the difference between upstream and downstream pressure. To calculate *β*, the inner diameter of the tube was measured, and the diameter of the 21-gauge needle (Becton, Dickinson and Company, Franklin Lakes, NJ) was determined via the Sigma-Aldrich Syringe Needle Gauge Chart.^41^ Finally, *ϵ* is the expansibility factor, which was calculated using data in CRANE Technical Paper 410M.^42^ The calculated mass flow rate was then converted to moles of nitrogen gas via equation (2).

$${mol}_{in} =\frac{Q_{m}*\Delta t}{{mw}_{N2}}$$

(2)

where *mol_in_* is the moles of nitrogen entering the pressure box, *Q_m_* is the mass flow rate, *∆t* is the difference between the current and previous time point, and *mw_N2_* is the molecular weight of the nitrogen gas. Moreover, the moles of gas already present in the container were calculated using the derivation of the specific ideal gas law below (3):

$${mol}_{cont}=\frac{V*P}{T*R_{u}}$$

(3)

where *mol_cont_* is the moles of gas within the container, *V* is the internal volume of the pressure chamber, *P* is the pressure at a given time point, *T* is the temperature, and *R_u_* is the universal gas constant. The outputs of equations 2 and 3 were combined, providing the total moles of gas present in the container after inlet of nitrogen. Because the volume of the pressure box remains fixed throughout the process, the initial and final conditions could be related as follows (4):

$$P_{cont}=\frac{\left( {mol}_{in}+{mol}_{cont} \right)}{mol_{cont}}P_{0}$$

(4)

where *P_cont_* is the final box pressure and *P_0_* is the initial pressure. For these calculations, it was assumed that the gaseous input did not cause a significant alteration to the volume of the box or its internal temperature. Because the flow rate is dependent on the pressure differential, the pressure was iteratively calculated every second to generate a curve (Figure 1D). The mathematical model illustrated that approximately 6 minutes of inlet time was necessary to equalize the container pressure with the regulator (225 kPa ≅ 2.25 atm).

The calculations were completed assuming ideal conditions; however, actual conditions offered stochastic factors, such as potential leakage at the needle/pipe interface. Therefore, the inflow time was lengthened to 10 minutes. The increase in inflow time did not pose a significant safety concern, as the mass flow rate exponentially decreased with the pressure differential.

#### Human Tissue Procurement

Patients provided written informed consent to collect and study their glioblastoma tissue in accordance with the study protocol submitted to and approved by St. Joseph’s Hospital and Medical Center Institutional Review Board. All patient data and tissue samples were de-identified to ensure anonymity. Tissue from six glioma patients was used in the experiments (Table S1). All the methods were performed in accordance with the relevant guidelines and regulations.

Upon collection, the brains were cut in 2-cm coronal slabs, which were submerged in a solution of 4% paraformaldehyde in PBS 0.1 M (i.e., PFA 4%) for 100 hours at 4°C. Tissue was dissected and preserved at 4°C in PBS 0.01% NaN_3_ for less than 2 years.^16^ For this study, 3 × 3 × 1-mm or, alternatively, anatomically adjacent samples were used in comparative experiments to minimize variability. Samples were obtained from tumor-free brain areas (i.e., cerebellum, cortex, or caudate nucleus), as ascertained by available imaging and gross inspection.

#### Cell Culture

All experimental protocols involving the use of human tissues were approved by the St. Joseph’s Hospital and Medical Center Institutional Review Board. All the methods were performed in accordance with the relevant guidelines and regulations.

A patient-derived glioblastoma cell line (GB3) was established from resected primary glioblastoma multiforme tumor tissue at Barrow Neurological Institute. Briefly, tumor tissue was processed using the gentleMACS Dissociator and Tumor Tissue Dissociation Kit (Miltenyi Biotec, Inc., Auburn, CA). Cells were expanded as neurospheres in tissue culture dishes coated with poly-2-hydroxyethyl methacrylate (Sigma-Aldrich, St. Louis, MO) or grown adherent on laminin (Thermo Fisher Scientific, Waltham, MA) in neural stem cell medium consisting of Dulbecco’s Modified Eagle Medium and F12-Glutamax, supplemented with B27 and N2 (Fisher Scientific, Hampton, NH), in the presence of 20 ng/mL epidermal growth factor and 20 ng/mL fibroblast growth factor-2 (MilliporeSigma, Burlington, MA). To generate the GB3–red fluorescent protein (GB3-RFP) cell line, we transduced GB3 cells with premade lentiviral particles (AMSBIO, Cambridge, MA) expressing RFP-luciferase, and selected them using blasticidin (2 µg/mL).

#### Mouse Glioblastoma Multiforme Xenografts

All animal procedures were performed following the guidelines of the Public Health Service Policy on Humane Care of Laboratory Animals and approved by the Institutional Animal Care and Use Committee of St. Joseph’s Hospital and Medical Center. Six-week-old *IcrTac:ICR-Prkdcscid* mice were used for in vivo orthotopic transplant of fluorescently tagged GB3-RFP cells. For orthotopic transplants, 2 μL of dissociated cells at a density of 100,000 cells/μL were injected in the right hemisphere (stereotaxic coordinates anteroposterior-0, mediolateral-2, dorsoventral-2.5), as described elsewhere.^43^ Four weeks after injection, tumor-bearing mice were euthanized with a lethal intraperitoneal injection of 2.5% Avertin (2,2,2-Tribromoethanol, Sigma-Aldrich, cat. no. T48402; *tert*-Amyl alcohol, Sigma-Aldrich, cat. no. A1685). Tissues were fixed through intracardial perfusion with Ringer solution (Electron Microscopy Sciences, cat. no. 11763-10) supplemented with 40 mM NaNO_2_, 2 mM NaCHO_3_, and 50 IU/mL heparin, followed by ice-cold 4% PFA in 0.1 M phosphate buffer (PB). Brains were subsequently cryoprotected with incubation in 30% sucrose PB for 48 hours before being cut into 1-mm coronal sections using a vibratome (Microm HM550, Thermo Fisher Scientific).^44^ For thin section stainings, brains were frozen at -50°C before being cut in coronal free-floating 40-µm–thick sections. Sections were preserved at -20°C in antifreezing solution until starting the staining procedure.

##### PACT Clearing

##### **Solution Preparation**

The PACT clearing procedure was applied as previously described^17,18^ with temperature modifications.^19^ Hydrogel solution was prepared by mixing 40 mL of 40% Acrylamide (Bio-Rad Laboratories, Inc., cat. no. 161-040), 1 g of VA-044 Initiator (Wako Chemicals USA, Inc., cat. no. 27776-21-2) in 360 mL of PBS. Clearing solution for human tissue was prepared by mixing 400 mL of 20% sodium dodecyl sulfate (Thermo Fisher Scientific, cat. no. 28365) and 200 mL of 1M boric acid (Sigma-Aldrich, cat. no. B7901) in 400 mL of distilled water, for a final concentration of 8% SDS. For mouse samples, a 4% SDS solution was used to maintain tissue integrity.

##### **Hydrogel Infusion and Washing**

The tissue sample was completely submerged in hydrogel solution in a 15-mL tube (Corning Falcon, Fisher Scientific) for 1 to 4 days at 4°C with gentle shaking and addition of fresh hydrogel solution every 2 days. Each tube was then degassed on ice for 10 minutes using a vacuum to remove all the O_2_, followed by an injection of N_2_ for 5 minutes. After 4 hours at room temperature, the tissue was transferred from the tube to the 12-well plate fitting the pressure box. Tissue was incubated in clearing solution at 40°C under continuous rocking, replacing the solution every 2 days. Upon completion of the clearing, the tissue was washed with boric acid buffer 0.2 M/0.1% Triton X at pH 8.5 for 2 days. After the washes were completed, we proceeded with the IHC protocol (see below).^18^

#### CUBIC Clearing

##### **Solution Preparation**

The CUBIC method was applied with slight modifications^19^ from the original protocol^12^ Reagent 1 (R1) was prepared by adding 30 g of urea (Sigma-Aldrich, cat. no. U0631), 30 mL of Quadrol (Sigma-Aldrich, cat. no. 122262), and 17 mL of Triton X-100 (Sigma-Aldrich, cat. no. T8532) to 42 mL of distilled water. R1 was then diluted 1:1 with water to generate water-diluted reagent 1. Reagent 2 (R2) was prepared by adding 31.6 g of urea (Sigma-Aldrich, cat. no. U0631), 52.4 g of sucrose (Sigma-Aldrich, cat. no. S0389), and 15 mL of triethanolamine (Sigma-Aldrich, cat. no. 90279) to 25 mL of distilled water. R2 was diluted 1:1 with 0.1 M PB to form PB-diluted reagent 2.^12^

Reagent 1

Every step of the procedure was performed in the 12-well plate under continuous rocking. After a wash in PB/0.01% NaN_3_ for 2 hours at room temperature, the tissue was incubated in water-diluted reagent 1 at 40°C for 5 hours, followed by washes in R1 solution for 6 days at room temperature, replacing the R1 every second day. On day 7, the sample was washed in PB/0.01% NaN_3_ for 2 hours at room temperature before commencing the IHC procedure.

##### Reagent 2

At the end of the IHC staining, the tissue was incubated with PB-diluted reagent 2 for 6 hours at room temperature. Afterward, the sample was incubated with R2 for 12 hours at 40°C. These last two steps were repeated once before mounting.

**iDISCO Clearing**

***Sample Pretreatment***

The iDISCO+ protocol was used without modification.^45^ The tissue was dehydrated via serial incubations in increasing concentrations of methanol (VWR cat. no. BDH2029) in H_2_O (20%/80%, 40%/60%, 60%/40%, 80%/20%, 100%/0%, 100%/0%), each for 1 hour at room temperature. The sample was then chilled at 4°C before being placed in a 66% dichloromethane (Sigma-Aldrich, cat. no. 270997)/ 33% methanol solution at room temperature overnight. After two washes in 100% methanol room temperature, the tissue was bleached in 5% H_2_O_2_ (Fisher Scientific, cat. no. H325-500)/95% methanol overnight at 4°C. The sample was then rehydrated in a methanol/H_2_O series (80%/20%, 60%/40%, 40%/60%, 20%/80%, 0.1M PB) for 1 hour each at room temperature before beginning the immunostaining procedure.

***Sample Clearing***

After the immunostaining, the tissue was dehydrated in methanol/H_2_O series (20%/80%, 40%/60%, 60%/40%, 80%/20%, 100%/0%, 100%/0%) for 1 hour each at room temperature. Subsequently, the sample was incubated in 66% dichloromethane/33% methanol solution at room temperature for 3 hours. The tissue was then washed in 100% dichloromethane for 15 minutes twice at room temperature. Finally, the sample was incubated in dibenzyl ether (Sigma-Aldrich, cat. no. 108014) until mounting.

#### IHC, pIHC, and Stainings

##### **Cleared Samples**

Timing of immunostaining in relation to the clearing procedure is depicted in Figure 1E. First, samples were repeatedly washed with PB/0.1% Triton-X (PBTX) before incubation with the primary antibodies for 72 hours at 4°C in PBTX/2% goat serum. See Table S2 for a comprehensive list of antibodies used in the study.

Following repeated washes in PBTX, samples were incubated with the species-matching Alexa-conjugated secondary antibodies (Table S3) at a dilution of 1:200 (Invitrogen, Carlsbad, CA) and 10 µg/mL DAPI in PBTX/2% goat serum for 72 hours at 4°C. Tomato lectin was added at a dilution of 1:250 (Vector Laboratories, Burlingame, CA), together with the secondary antibodies. Negative controls were subjected to the same procedure (two 72-h pressurizations) without adding the primary antibody. We calculated that the changes in temperature during protocol would result in negligible changes in pressure.

##### **Thin Sections**

The procedures are schematically represented in Figure 7A.

*Conventional IHC (19 h):* After repeated washing in PB, free-floating 40-µm–thick mouse brain sections were blocked with PBTX/10% goat serum for 2 hours at room temperature. Primary antibodies were incubated for 11 hours at 4°C in PBTX/2% goat serum. After repeated washes, sections were incubated with species-matching Alexa-conjugated secondary antibodies at a dilution of 1:1000 (Invitrogen) for 2 hours at 4°C in PBTX/2% goat serum.

*pIHC (8 h) and control IHC (8 h):* The blocking, primary and secondary antibody steps were performed as described above, changing the length of each step to 30 minutes, 2 hours and 1 hour, respectively.

Antibody concentrations are summarized in Tables S2 and S3. DAPI incubation was performed in PB for 10 minutes. Sections were mounted on slides using ProLong Gold Antifade mountant (Invitrogen).

#### Thick Specimen Mounting

As described by Treweek et. al., RIMS was made by adding 40 g of Histodenz (Sigma-Aldrich, cat. no. D2158) to 30 mL of 0.02 M PB/0.01% NaN_3_.^17^ Immunostained samples, either PACT-cleared or CUBIC-cleared, were incubated in RIMS solution for 6 hours. Blu Tack (Bostik, Milwaukee, WI) was used on conventional glass slides (VWR International) to create a 1.5-mm–thick chamber where a single sample was submerged in RIMS and covered with a 0.15-mm coverslip. Slides were stored at 4°C. For iDISCO-cleared tissue, dibenzyl ether was used as the mounting medium.

Spectrophotometry—Measurement of Collimated Light Transmittance

After RIMS incubation, light transmittance (400–800 nm, with 10 nm steps) of the 1-mm–thick human brain tissue blocks were measured with a FlexStation 3 spectrophotometer (Molecular Devices, San Jose, CA).

#### Electron Microscopy

Control, cleared, and cleared-pressurized human brain samples were post-fixed in 2% osmium tetroxide, dehydrated, and embedded in Durcupan resin (Fluka; Sigma-Aldrich). Semithin sections (1.5 mm) were cut with a diamond knife and stained with 1% toluidine blue for light microscopy. Ultrathin sections (70–80 nm) were cut, stained with lead citrate, and examined under an FEI Tecnai G^2^ Spirit transmission electron microscope (FEI Europe B.V., Eindhoven, Netherlands) using a digital camera (Morada Soft Imaging System, Olympus, Tokyo, Japan). The analysis was performed blindly.

**Tissue Deformation Analysis**

The procedure for macroscopic analysis of tissue deformation was adopted from Neckel et al., 2016^46^ and Wan et al., 2018.^2^ Briefly, 1-mm thick sections obtained from the striatum of 3 different human brains were cleared according to the CUBIC, PACT and iDISCO protocols described above. The individual samples were photographed before clearing and after the key steps of each procedure. The surface area of each tissue sample was determined using ImageJ (public domain, National Institutes of Health). For each technique (iDISCO, CUBIC, PACT) the difference in surface area change between the IHC and pIHC groups was compared.

#### Imaging and Analysis

Cleared tissues were imaged using the Leica SPE system utilizing a 10× dry objective (numerical aperture of 0.30). Alternatively, image data were collected using a Leica TCS SP8 LSCM [NIH SIG award 1 S10 OD023691-01] housed in the W.M. Keck Bioimaging Facility at Arizona State University using a 25× water objective (numerical aperture of 0.85) (Leica Camera, Wetzlar, Germany).

Comparative analysis of fluorescence intensity was performed on 400-µm confocal stacks (10-µm interval, thick tissue). All comparative imaging was taken with identical parameters, which were set on the most intense superficial signal, using the look-up table Leica feature. Thick tissue was imaged at the center, avoiding the sides, which would introduce bias because of lateral antibody penetration. In line with previous work,^7,47,48^ confocal stacks are shown in figures as 3-D images created using the 45° projection visualization on Imaris (Bitplane, Belfast, Ireland). Thin sections were imaged using a 20× oil objective using 40-µm stacks (1-µm interval). Image analysis was performed with Imaris and ImageJ. Experiments were repeated three or more times. Graphs show data averaged from repeated measurements of at least three independent samples. Fluorescence intensity was normalized for each staining on the highest value obtained in each experiment. Data plotting and statistical analysis were performed with GraphPad Prism (GraphPad Software, La Jolla, CA). Statistical analysis was done through two-way analysis of variance (alpha=0.05) with the Šidák method for multiple comparisons.
