## Supplementary material for "Enhanced tissue penetration of antibodies through pressurized immunohistochemistry": Figure Legends S1-S6

**SUPPLEMENTAL FIGURE LEGENDS**

**Figure S1.** Set-up of pressurization device. **(A)** Photograph of components required for pressure chamber construction. **(B)** Photograph of pressure device, sealed with parafilm before pressurization. **(C)** Photograph of fastened pressure device during pressurization. Box shows regulator pressure gauge reading during nitrogen inlet. *Used with permission from Barrow Neurological Institute, Phoenix, Arizona.*

**Figure S2.** pIHC does not affect antibody penetration in iDISCO-cleared human tissue. **(A)** Graph shows the mean relative surface area of human striatal brain tissue samples after iDISCO clearing and IHC with and without pressurization. iDISCO induces a 25% tissue shrinkage, with no differences between conditions. Error bars show SE. N=3 independent samples per condition. Photographs depict human striatal brain tissue sections before (top) and after (bottom) the iDISCO protocol. **(B, C)** 400-μm thick acquisition of iDISCO-cleared samples immunostained for IBA1 **(B**, *red***)**, α-SMA **(B**, *green***),** GFAP **(C**, *green***)**, NeuN **(C**, *red***)**, and lectin-649 **(C**, *blue***)**, using free diffusion (top rows) or pIHC (bottom rows). Both pIHC and free diffusion were unable to effectively penetrate and label the superficial 400 μm of iDISCO-cleared tissue, regardless of the antibody. **(D)** Snapshots of the surface stainings for IBA1 (top, *red*), NeuN (middle, *red*), GFAP (bottom, *green*), and lectin (bottom, *blue*) demonstrate the morphological specificity of each staining, despite poor penetration. Scale bars in **B**, **C**, 200 μm; in **D**, 50 μm (all panels). All experiments were independently performed three or more times; representative images are shown. *Used with permission from Barrow Neurological Institute, Phoenix, Arizona.*

**Figure S3.** 3-D labeling of vascular and neuronal markers in human brain samples using pIHC. **(A and B)** Isometric **(A)** and top **(B)** views of 800-μm thick CUBIC-cleared human striatal tissue sample. pIHC allows for uniform CD31 (*red*) staining. Image shows a z-stitched composite of 400-μm confocal acquisitions. **(C)** Image shows a 2.1 × 1.4-mm*,* 750-μm–thick mosaic acquisition of a human cerebellar tissue sample cleared with PACT and immunostained with pIHC for neurofilament (*green*). Although the low-density molecular layer (ML) reveals the fine details of neurofilament+ Purkinje cells, the high cell density of the granule layer (GL) impedes antibody penetration below 120 μm from the surface (*arrows*). Signal saturation in GL is due to overlapping of the intense neurofilament signal in the first 120 μm of depth. (**D**) Close-up 2-D image of superficial neurofilament immunostaining (z=40 μm) showing the bona fide morphological features of Purkinje cells and the neurite network. **(E, F)** Comparison of the intensity and depth of an anti-NeuN (*red*) antibody tested on CUBIC-cleared human hippocampus tissue samples. Image in **E** shows an 830-μm thick mosaic acquisition of the dentate gyrus (x, y = 2.4 × 1.9 mm). Image in **F** shows a 930-µm acquisition (x, y = 1.1 × 1.1 mm). Both isometric (top row) and top views (bottom row) of the dentate gyrus illustrate the improved brightness and depth associated with pIHC relative to free diffusion. **(G)** Graph shows quantification of normalized mean intensity measured along the z-axis at constant laser intensity comparing pIHC and IHC (Ctrl) for NeuN. Pressurization (*black*) significantly improves staining intensity compared to free diffusion (*gray*). Error bars show SE. N=3 independent samples. Scale bars in **A,** 200 μm; in **B**, 100 μm; in **C,** 500 μm; in **D**, 100 μm; in **E** and **F**, 500 μm. All experiments were independently performed three or more times; representative images are shown. *Used with permission from Barrow Neurological Institute, Phoenix, Arizona.*

**Figure S4.** Analysis of whole-tissue structural deformation. **(A)** Images of human brain tissue during different stages of the CUBIC clearing protocol. **(B)** Graph shows the relative surface area of the human samples after each step in the CUBIC procedure. All measurements were normalized to the surface area of the untreated samples (dotted line). Following a first increase after the R1 solution, immersion in the aqueous solutions during the IHC procedure results in a transient decrease in size. Another significant swelling occurs after incubation in R2 solution. Tissue shrank again in the final incubation in RIMS, with a final size approximately 25% larger than the untreated samples. Pressurization (*grey* bars in graphs) did not induce any difference compared to control at any stage of the protocol. **(C)** Images of human brain tissue during stages of PACT clearing. **(D)** Graph demonstrates the relative surface area of the human samples after each step in PACT. Like CUBIC-cleared tissue, PACT-cleared tissue shows dynamic changes in size with the different steps of the procedure, with a first significant swelling during the incubation in SDS. Pressurization did not influence the variations in tissue size compared to control. A 50% increase in size is finally measured when incubated with RIMS before mounting. Bars in graphs show N=3 independent samples of human striatal tissue; representative images are shown. Error bars show SE. Scale bars in **A**, **C**: 500 µm. *Used with permission from Barrow Neurological Institute, Phoenix, Arizona.*

**Figure S5.** Vascular-associated artifacts. **(A)** A 3-D view of 400-μm confocal acquisition showing negative control experiments. Background fluorescence intensity is decreased in pressurized samples (bottom row) compared with controls (top row). PACT samples (right column) show co-occurrence of vascular-associated autofluorescence in the 488-nm and 568-nm excitation wavelengths, which is absent in CUBIC (left column) samples. **(B)** TEM analysis showing a lack of identifiable erythrocytes (er) in CUBIC samples. The *red* contour shows blood vessel basal lamina. **(C)** Images of human striatum immunostained with an anti-laminin antibody, showing nonspecific granulation on the surface in PACT-cleared but not in CUBIC-cleared tissue. Scale bars in **A,** 200 μm; in **B**, 2 μm; in **C**, 100 μm. All experiments were independently performed three or more times; representative images are shown. *Figures S5A and S5C are used with permission from Barrow Neurological Institute, Phoenix, Arizona. Figure S5B is used with permission from University of València, València, Spain.*

**Figure S6.** 3-D reconstruction of the neurogenic niche. **(A)** 2-D image of migrating neuroblasts pIHC-stained with anti-doublecortin (DCX) in the anterior forebrain neurogenic areas cleared with CUBIC in control mice, demonstrating bona fide morphological features of DCX. **(B)** Mosaic confocal acquisition demonstrating DCX labeling specificity in the subventricular zone (SVZ). **(C)** Composite images showing sequential scans of the transition area between SVZ (right) and the rostral migratory stream (RMS, left). The right image is a mosaic of nine acquisitions shown with 30° tilting along the Y axis to show the SVZ and simultaneously emphasize the full penetration of the DCX staining in the complex 3-D arrangement of this structure. The RMS in the left image is shown with a 90° angle with respect to the viewer. DCX stainings were independently performed three times; representative images are shown. Scale bar in **A**, 50 μm; in **B**, 500 μm; in **C**, 200 μm. *Used with permission from Barrow Neurological Institute, Phoenix, Arizona.*
