## Supplementary figures and images for "Enhanced tissue penetration of antibodies through pressurized immunohistochemistry"

### Fig S1

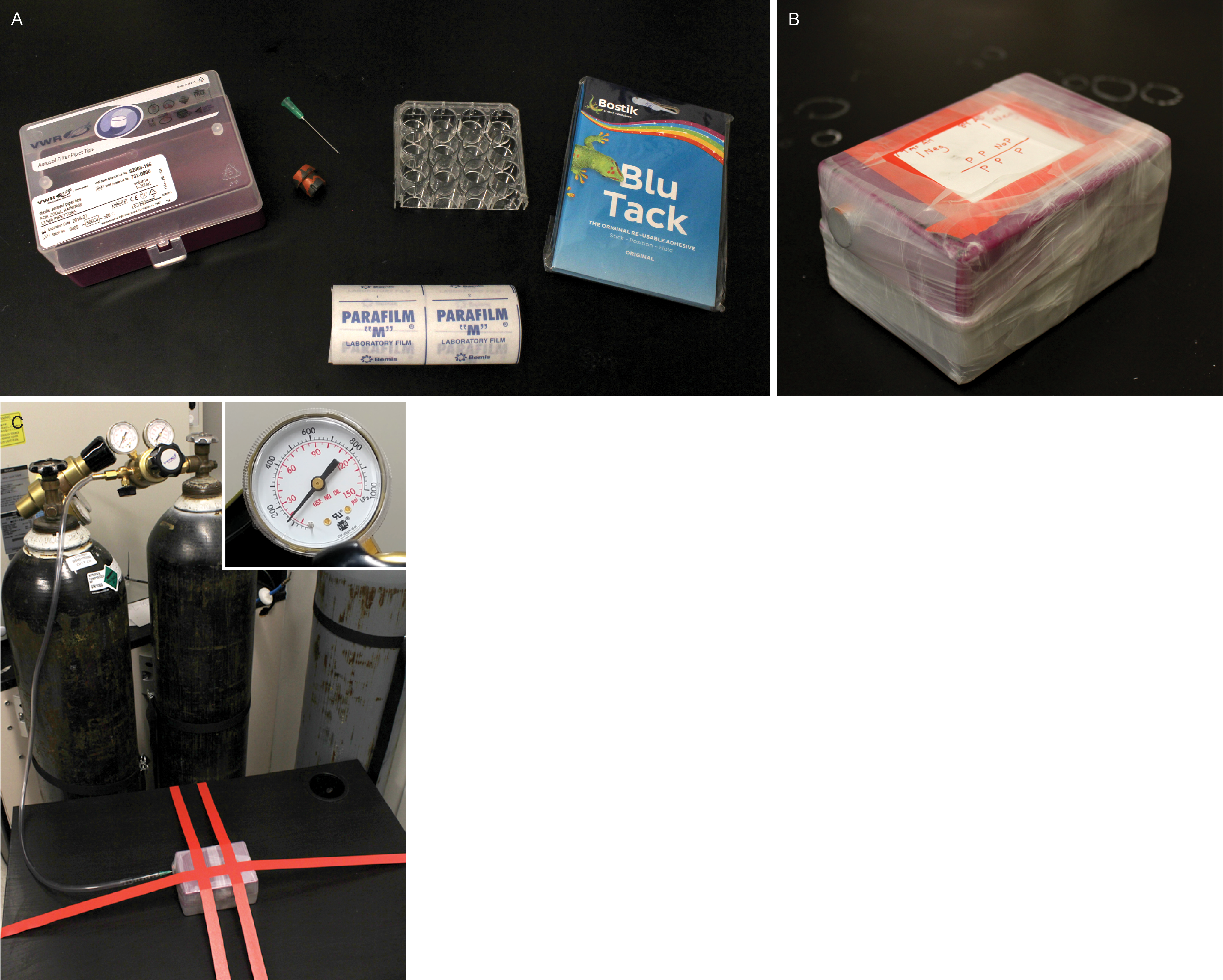

### Fig S2

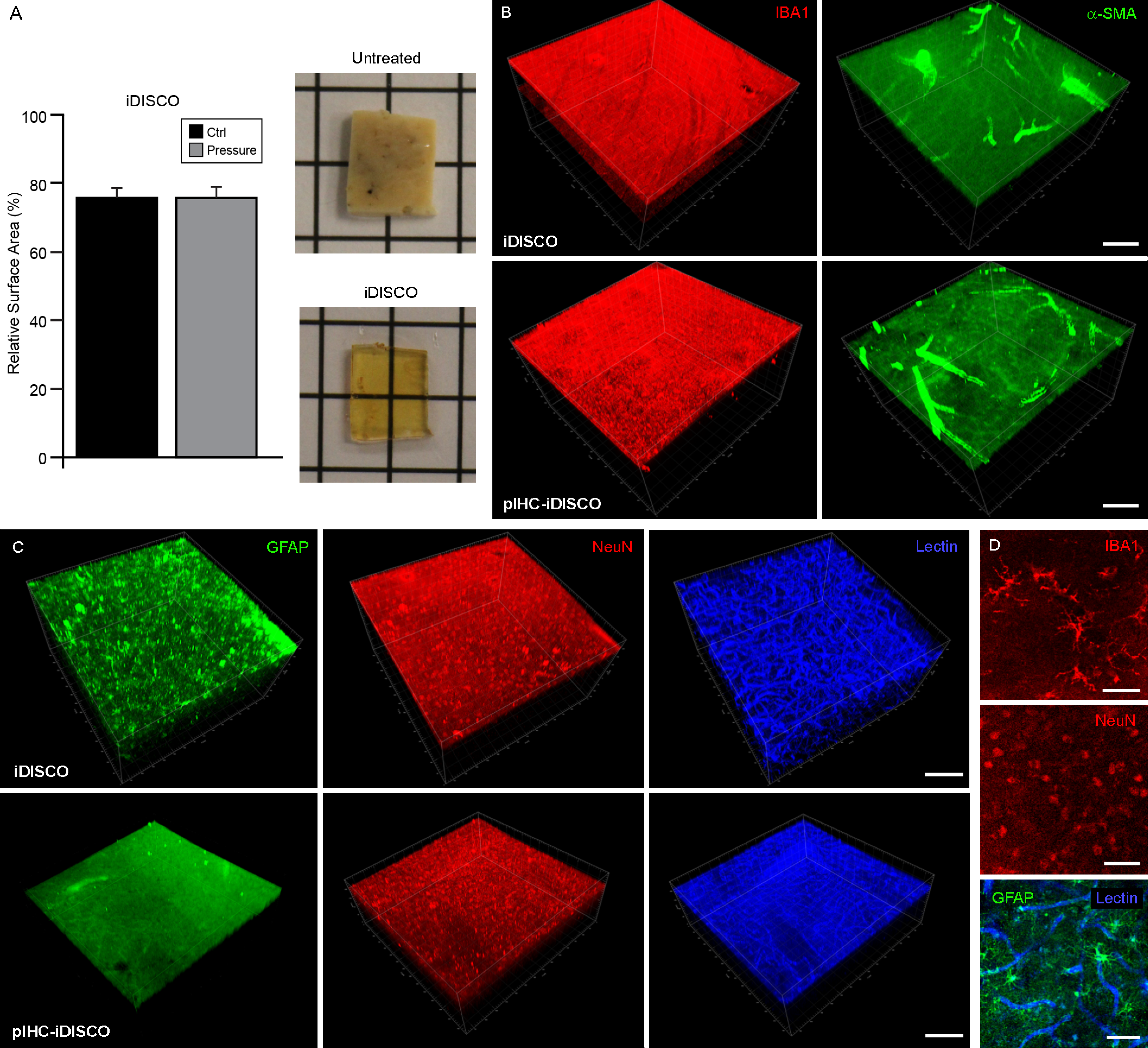

### Fig S3

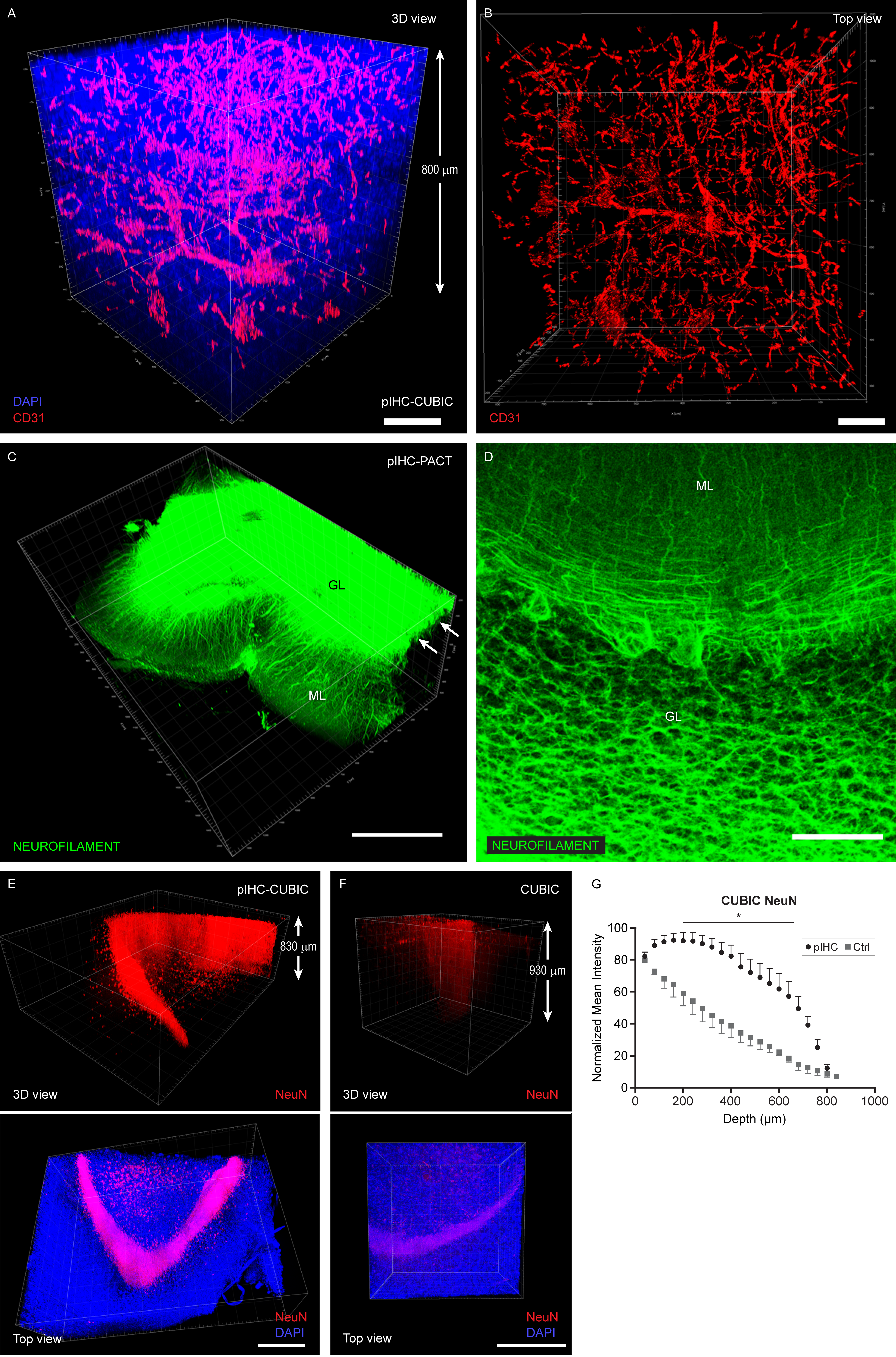

### Fig S4

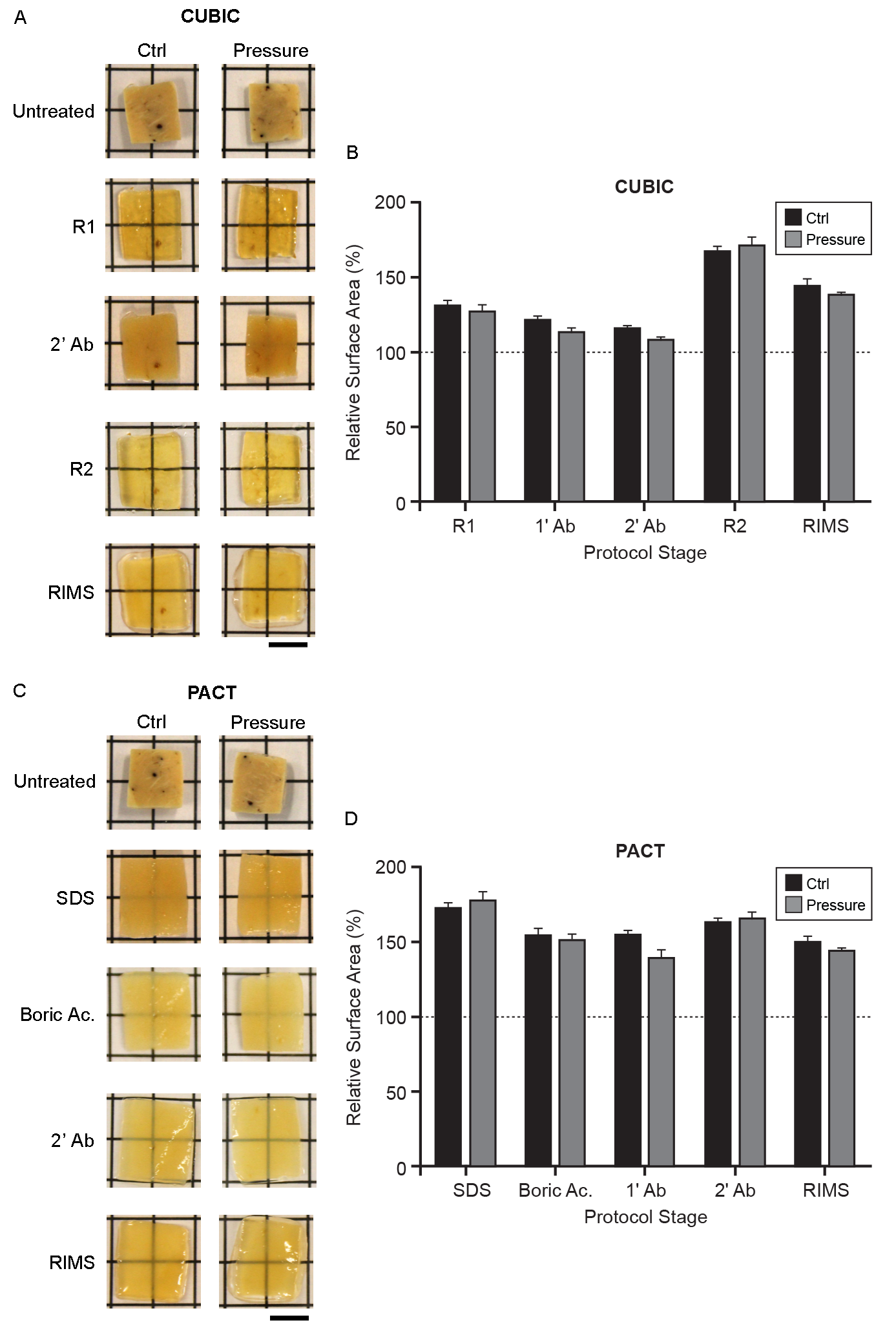

### Fig S5

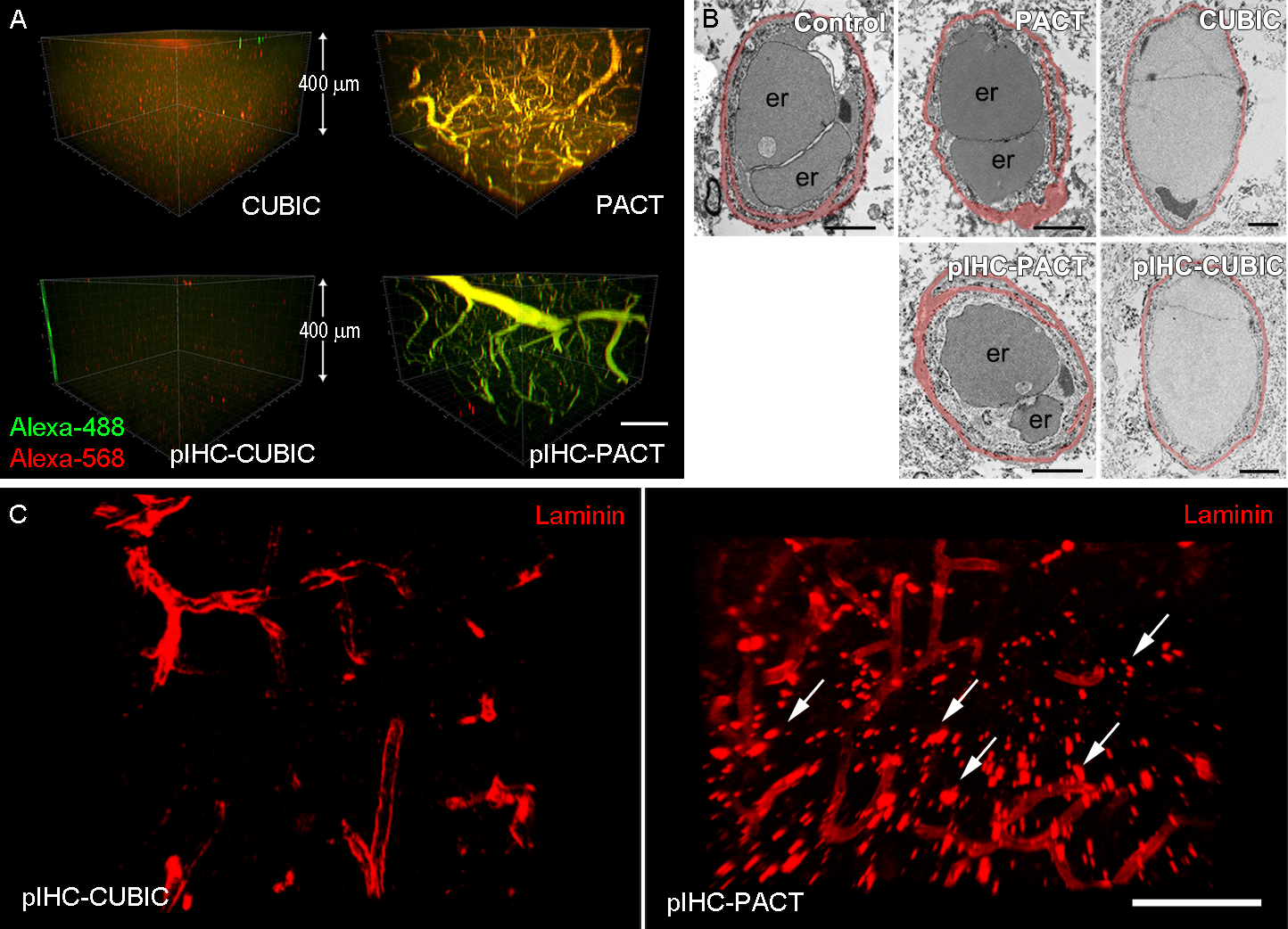

### Fig S6

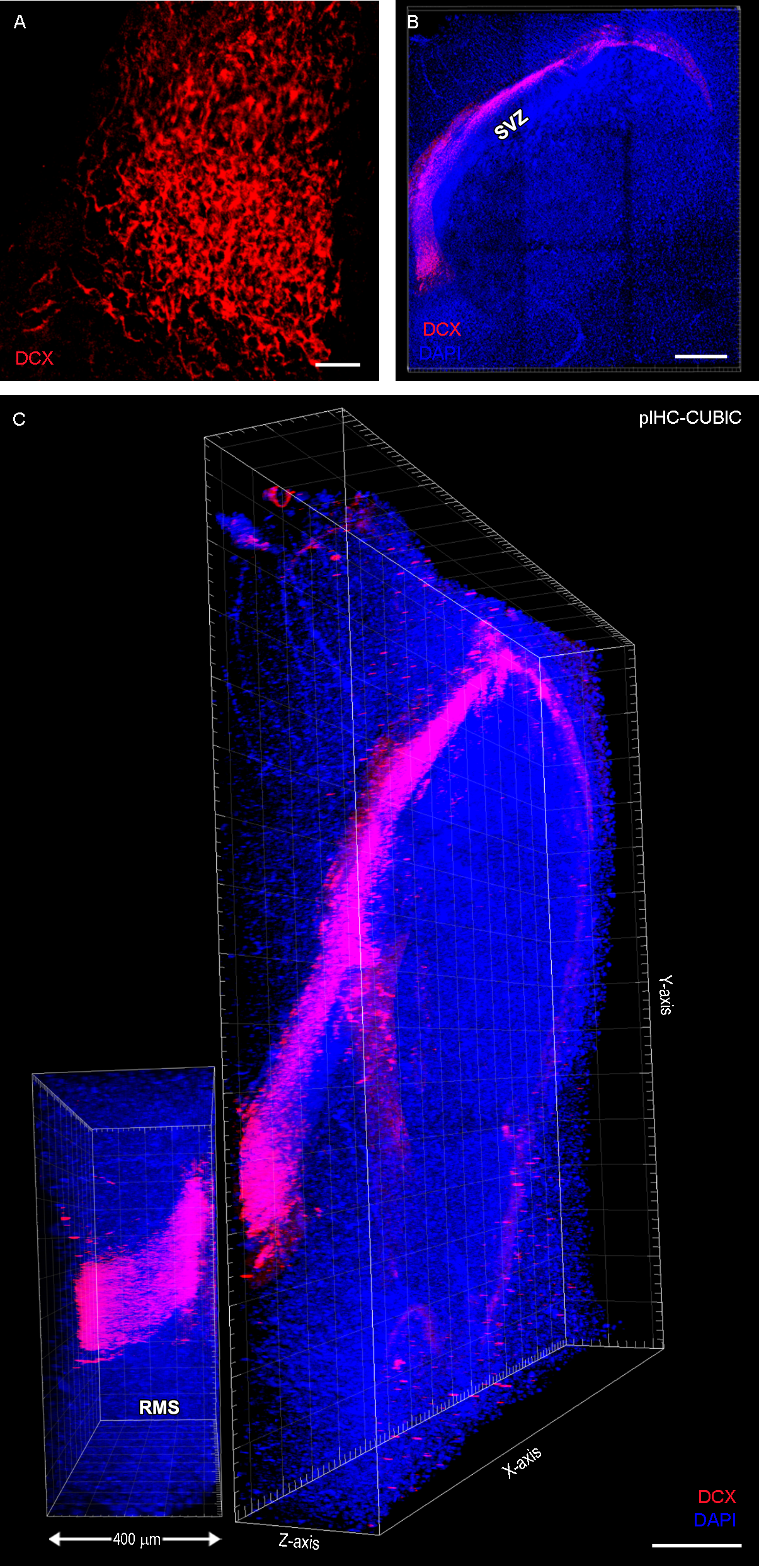
