## Supplementary material for "Enhanced tissue penetration of antibodies through pressurized immunohistochemistry": Video Legends S1-S4

**SUPPLEMENTAL VIDEO LEGENDS**

**Video S1.** Narrated practical demonstration of 1) building pressure chamber, 2) pressurizing the chamber, and 3) mounting samples. *Used with permission from Barrow Neurological Institute, Phoenix, Arizona.*

**Video S2.** 3-D projection of a confocal acquisition of IBA1+ microglia on pIHC-CUBIC stained human caudate. Dimensions (x, y, z): 734 × 734 × 960 μm, z-stack size 10 μm. *Used with permission from Barrow Neurological Institute, Phoenix, Arizona.*

**Video S3.** 3-D projection of a confocal acquisition of GFAP+ astrocytes on pIHC-CUBIC stained human caudate. Dimensions (x, y, z): 230 × 230 × 510 μm, z-stack size 10 μm. *Used with permission from Barrow Neurological Institute, Phoenix, Arizona.*

**Video S4.** 3-D projection of a confocal acquisition of tomato-lectin+ vasculature (*cyan*), human xenografted RFP+ glioblastoma cells (*red*), and the proliferation marker Ki-67 (*green*) on pIHC-CUBIC stained mouse brain. Dimensions (x, y, z): 1030 × 1880 × 470 μm, z-stack size 10 μm. *Used with permission from Barrow Neurological Institute, Phoenix, Arizona.*
