## Supplementary material for "Enhanced tissue penetration of antibodies through pressurized immunohistochemistry": Tables S1-S3

**SUPPLEMENTAL TABLES**

**Table S1.** Human samples used in the study*

| **ID** | **Age, years** | **Sex** | **PMI (h)** |
| --- | --- | --- | --- |
| 0907 | 70 | F | 12 |
| 0209 | 64 | M | 14 |
| 1908 | 27 | M | 7 |
| 0611 | 20 | F | 13 |
| 1108 | 82 | M | 6 |
| 0926 | 73 | F | 4 |

*Cause of death for all patients was glioblastoma.

Abbreviations: PMI, postmortem interval.

**Table S2.** Specifics of primary antibodies and dyes tested in the study

| **Species** | **Antigen** | **Company** | **Cat. No.** | **Dilution (40 μm)** | **Dilution (1 mm)** |
| --- | --- | --- | --- | --- | --- |
| Chicken | Vimentin | Millipore | AB5733 | 1:1000 | 1:200 |
| Mouse | Fibronectin | Abcam | ab26245* | 1:200 | 1:50 |
| Rabbit | Laminin | Abcam | ab11575 | 1:200 | 1:50 |
| Guinea pig | MAP2 | Synaptic Systems | 188004 | 1:1000 | 1:100 |
| Rabbit | IBA1 | Wako | 019-19741 | 1:400 | 1:100 |
| Mouse | GFAP | Millipore | MAB360 | 1:500 | 1:50 |
| Mouse | α-SMA | Abcam | ab7817 | 1:400 | 1:50 |
| Mouse | OLIG2 | Millipore | AB9610 | 1:200 | NA |
| Mouse | Neurofilament | Abcam | ab7794 | 1:200 | 1:50 |
| Rabbit | RFP | Abcam | ab62341 | 1:500 | 1:50 |
| Mouse | NeuN | Millipore | MAB377 | 1:200 | NA |
| Rabbit | NeuN | Cell Signaling | 24307S | 1:200 | 1:25 |
| Rabbit | SOX2 | Cell Signaling | 3579S | 1:200 | 1:50 |
| Mouse | Ki-67 | DAKO | M7240 | 1:150 | 1:50 |
| Rabbit | Doublecortin | Cell Signaling | 4604S | 1:250 | 1:50 |
| Mouse | SMI-32 | Millipore | 559844* | 1:250 | 1:50 |
| Rabbit | CD31 | Abcam | ab28364 | 1:100 | 1:25 |
| - | DAPI | Invitrogen | d21490 | 1 μg/μL | 10 μg/μL |

Abbreviations: α-SMA, alpha–smooth muscle actin; DAPI, 4′,6-diamidino-2-phenylindole; GFAP, glial fibrillary associated protein; IBA1, ionized calcium-binding adapter molecule 1; MAP2, microtubule associated protein 2; NeuN, neuronal nuclear antigen; OLIG2, oligodendrocyte transcription factor 2; RFP, red fluorescent protein; SOX2, sex determining region Y-box 2; SMI-32, neurofilament H non-phosphorylated.

*Discontinued.

**Table S3.** Specifics of secondary antibodies and dyes tested in the study

| **Target** | **Species** | **Company** | **Cat. No.** | **Wave­length (nm)** | **Dilution (40 μm)** | **Dilution (1 mm)** |
| --- | --- | --- | --- | --- | --- | --- |
| Mouse | Goat | Invitrogen | A11001 | 488 | 1:1000 | 1:200 |
| Mouse IgG1 | Goat | Invitrogen | A21121 | 488 | 1:1000 | 1:200 |
| Rabbit | Goat | Invitrogen | A11011 | 568 | 1:1000 | 1:200 |
| Rabbit | Goat | Invitrogen | A21245 | 647 | 1:1000 | 1:200 |
| Chicken | Goat | Invitrogen | A21449 | 647 | 1:1000 | 1:200 |
| Mouse | Goat | Jackson | 115-065-003 | Biotin | 1:1000 | 1:200 |
| Guinea pig | Goat | Invitrogen | A11075 | 568 | 1:1000 | 1:200 |
| Lectin |  | Vector | DL-1177 | 488 | 1:1000 | 1:250 |
| Lectin |  | Vector | DL-1178 | 647 | 1:1000 | 1:250 |
| Streptavidin |  | Invitrogen | S11226 | 568 | 1:500 | 1:100 |
