## Supplementary material for "Enhanced tissue penetration of antibodies through pressurized immunohistochemistry": Permission letter - Fig 1, 2, 3A-E, 4, 5A, 6, 7, S1-S4, S5A, S5C, S6, and Vid S1-S4

BARROW Neurological Institute® | St. Joseph's Hospital and Medical Center A Dignity Health Member

Barrow Neurological Institute holds the copyright to Figures 1, 2, 3A-E, 4, 5A, 6, 7, S1-S4, S5A, S5C, S6, and Videos S1-S4 to be used in the manuscript by Fiorelli R, Sidhu GS, Cebrian-Silla A, Melendez EL, Mehta S, Garcia-Verdugo JM, and Sanai N entitled "Enhanced tissue penetration of antibodies through pressurized immunohistochemistry" for possible publication in *bioRxiv*. Permission is granted for print publication and corresponding electronic publication of the manuscript and for all compilations that include this manuscript in its original form. Permission is granted with distribution rights in all languages throughout the world. The permission is subject to the use of a standard credit line on the same page where our figures and/or videos will appear. Permission is also granted to use the figures for the Journal cover or promotion of the issue or Journal.

**PLEASE NOTE:** Permission to use this material in other forms and for other uses requires a separate permission request. Permission to distribute or sell the figures and/or videos independent of the original manuscript and corresponding electronic publication of the manuscript requires a separate permission request. When submitting a request for other uses, please return a copy of this approved permission request to use this material together with the name, anticipated publication date, and tentative price of the other format.

Cassie Todd [signature] Date: 8/18/2020

**Cassie Todd, Production Editor**
Neuroscience Publications
Barrow Neurological Institute

350 W. Thomas Rd.
Phoenix, AZ 85013
602.406.3593
barrowneuro.org
