## Supplementary material for "Enhanced tissue penetration of antibodies through pressurized immunohistochemistry": Permission letter - Fig 3F, 5B-E, S5B

BARROW
Neurological Institute®

St. Joseph's Hospital
and Medical Center
A Dignity Health Member

University of Valencia holds the copyright to IM 20161-01, 20168-01, 20169-01, and 20171-01 – 20173-01 to be used in the manuscript by Fiorelli R, Sidhu GS, Cebrian-Silla A, Melendez EL, Mehta S, Garcia-Verdugo JM, and Sanai N entitled "Enhanced tissue penetration of antibodies through pressurized immunohistochemistry" for possible publication in *bioRxiv*. Permission is granted for print publication and corresponding electronic publication of the manuscript and for all compilations that include this manuscript in its original form. Permission is granted with distribution rights in all languages throughout the world. The permission is subject to the use of a standard credit line on the same page where our figures and/or videos will appear. Permission is also granted to use the figures for the Journal cover or promotion of the issue or Journal.

**"Used with permission from University of Valencia."**

**PLEASE NOTE:** Permission to use this material in other forms and for other uses requires a separate permission request. Permission to distribute or sell the figures and/or videos independent of the original manuscript and corresponding electronic publication of the manuscript requires a separate permission request. When submitting a request for other uses, please return a copy of this approved permission request to use this material together with the name, anticipated publication date, and tentative price of the other format.

Date: May 22nd, 2020

**Jose M. Garcia-Verdugo, PhD**
University of Valencia
Avda. Vicente Andrés Estellés s/n
C.P.: 46100
Burjassot Valencia, Spain
